# SatelliteQ: an open-source workflow for multidimensional, object-based quantification of centriolar satellites

**DOI:** 10.64898/2026.10.07.756960

**Authors:** Efe Begar, Ezgi Odabasi, Lucie Menguy, Sophie Saunier, Elif Nur Firat-Karalar

## Abstract

Centriolar satellites are membrane-less organelles that store, modify and traffic proteins, in particular those of the centrosome and the primary cilium, and their deregulation is linked to ciliopathies, neurodevelopmental and neurodegenerative disorders. Satellite number, size, intensity and spatial distribution vary with cell type and cell state, yet satellites are usually quantified by a single readout, the pericentrosomal fluorescence intensity in maximum-projected images, which cannot distinguish changes in satellite number, size and position. Here we present SatelliteQ, an open-source Fiji and Python workflow for object-based quantification of satellites in 2D, 3D and time-lapse images. SatelliteQ segments satellites, detects centrosomes or primary cilia in the same cells, and reports satellite number, size, intensity, distance to the centrosome, shape and spatial distribution per satellite and per cell, with interactive graph and spatial viewers. Its measurements correlate closely with Imaris-based quantification; absolute values depend on segmentation settings and analysis mode, whereas the biological differences tested here were preserved across matched settings. Using SatelliteQ, we resolved distinct spatial signatures of satellite dispersal induced by microtubule depolymerization and by disruption of weak hydrophobic interactions, quantified differences between cell types and the scaling of satellite number with cell size, captured PCM1 granule dynamics in living cells, and compared satellite organization between ciliated and unciliated cells. Depletion of the ciliopathy-associated satellite protein CEP290 altered satellite abundance and pericentrosomal organization, while patient-derived cells carrying CEP290 mutations displayed divergent satellite phenotypes that were not explained by ciliation alone. SatelliteQ thus establishes satellite properties as a multidimensional cellular phenotype and provides an accessible framework for exploring satellites as an organelle-level readout in disease.

## INTRODUCTION

Centriolar satellites (hereafter satellites) are membrane-less organelles first described by electron microscopy in epithelial cells as approximately 100 nm electron-dense granules that concentrate and move around the centrosome in a microtubule-dependent manner (Kubo et al., 1999). Their scaffolding protein, pericentriolar material-1 (PCM1), is essential for satellite assembly (Kubo et al., 1999; Dammermann and Merdes, 2002; Begar et al., 2026). Proteomic studies of PCM1 have identified over 200 associated proteins, referred to as satellite clients, which are strongly enriched in centrosome and primary cilium components (Aydin et al., 2020; Gheiratmand et al., 2019; Gupta et al., 2015; Quarantotti et al., 2019). The centrosome is the main microtubule-organizing center of animal cells, and in quiescent cells, its mother centriole templates the primary cilium, a microtubule-based cellular projection that transduces extracellular signals such as Hedgehog. By regulating the abundance, localization and modification of their clients, satellites contribute to cilium biogenesis and signaling, cell division, stress responses, proteostasis, neurogenesis and sperm maturation (Aydin et al., 2020; Begar et al., 2024; Hall et al., 2023; Odabasi et al., 2019; Wang et al., 2016; Zhao et al., 2021). Consistent with these functions, loss of PCM1 in mice causes developmental, psychomotor and cognitive abnormalities that mirror those of ciliopathies and schizophrenia, as well as male infertility (Hall et al., 2023; Huang et al., 2025; Monroe et al., 2020). In humans, PCM1 itself is linked to schizophrenia and, through gene fusions with tyrosine kinases, to leukemia, and satellite clients are mutated in ciliopathies, in neurodevelopmental disorders and in neurodegenerative diseases such as amyotrophic lateral sclerosis and Huntington’s disease (reviewed in Begar et al., 2024).

Consistent with their diverse functions, satellites exhibit substantial heterogeneity in their organization, both within individual cells and across cell types (Begar et al., 2026; Kubo and Tsukita, 2003; Odabasi et al., 2020). In most cultured mammalian cells, satellites are observed as sub-resolution granules that concentrate around the centrosome but are also distributed throughout the cytoplasm, with individual satellites differing in size and composition even within the same cell (Firat-Karalar et al., 2014; Gheiratmand et al., 2019; Kubo et al., 1999). Satellite properties are also dynamically remodeled with cell state. They disassemble at mitotic entry and reassemble after cell division (Dammermann and Merdes, 2002), reorganize in response to cellular stress (Tollenaere et al., 2015; Villumsen et al., 2013), change in spatial distribution and client composition during primary cilium formation (Kobayashi et al., 2014; Odabasi et al., 2023), and disperse upon microtubule depolymerization (Dammermann and Merdes, 2002; Kubo et al., 1999). Across cell types, satellites can also adopt distinct organizational states, ranging from micrometer-sized granules in multiciliated epithelial cells to a matrix surrounding the nuclear envelope in myotubes and liquid-like structures within the spindle domain of oocytes (So et al., 2019; Sorokin, 1968; Srsen et al., 2009; Zhao et al., 2021). Moreover, satellites assemble through a hierarchical pathway in which PCM1 multimerization generates a scaffold that recruits clients such as CEP290 in a defined order, and disruption of this pathway compromises ciliary signaling and mitotic progression (Begar et al., 2026). Together, these findings establish satellite properties as a multidimensional cellular phenotype that varies in number, size, morphology, abundance, and spatial distribution.

Despite this complexity, satellite phenotypes have generally been quantified using one or a limited number of readouts. In most studies, including our own, PCM1 or another satellite client is quantified as fluorescence intensity within a fixed-radius region around the centrosome in maximum-intensity projections (e.g., (Aydin et al., 2020; Kim et al., 2008; Odabasi et al., 2019; Villumsen et al., 2013)), or satellites are classified as clustered, dispersed, disassembled, dissolved, or absent. Intensity has also been measured in concentric rings around the centrosome to describe the radial distribution of satellites (Galati et al., 2018; Gheiratmand et al., 2019; McCurdy et al., 2022). These approaches capture major changes in pericentrosomal abundance or distribution but leave most satellite properties unresolved. Because the signal from all satellites within the analyzed region is summed, pericentrosomal intensity cannot distinguish changes in satellite number, size, or total PCM1 fluorescence intensity, and it does not capture cytoplasmic satellites or heterogeneity between individual satellites. Study-specific region sizes and maximum-intensity projections further limit comparability and can merge satellites at different depths. Direct counting has also been reported but remains uncommon. An early study estimated approximately 300 to 500 PCM1-positive granules per cell across several cultured cell lines, including epithelial cells and fibroblasts, and found that their abundance and degree of pericentrosomal clustering varied between cell types (Kubo and Tsukita, 2003). Manual counting is also subjective and impractical for large datasets. Because different perturbations and cell states can alter distinct combinations of satellite properties, resolving these properties separately is important for defining satellite phenotypes during biogenesis, maintenance and cellular perturbation.

In our recent work, object-based analysis of 3D images using commercial software enabled us to quantify satellite number, volume, total fluorescence intensity, mean fluorescence intensity per satellite, and distance from the centrosome (Begar et al., 2026). These measurements allowed us to distinguish changes in satellite number, size and PCM1 fluorescence during de novo assembly, characterize PCM1 truncation mutants with distinct granule properties, and follow client recruitment and remodeling during ciliogenesis. However, this approach requires a commercial license and custom pipelines that are rarely shared. For primary cilia, open-source tools such as CiliaQ, ACDC and detectCilia provide standardized morphological and intensity measurements in 2D and 3D, but no comparable open-source framework exists for centriolar satellites (Budde- Sagert et al., 2024; Hansen et al., 2021; Lauring et al., 2019). A standardized and accessible approach is therefore needed to resolve multidimensional satellite phenotypes and determine how individual properties change across cellular contexts and in response to perturbation and disease.

This gap is particularly relevant to ciliopathies, in which patient-derived cells are typically phenotyped based on cilium number, length and composition, increasingly with automated tools such as ALPACA, which quantifies ciliation, cilium length and intraflagellar transport in patient fibroblasts to support diagnosis (Doornbos et al., 2021). However, many ciliopathy-associated proteins, including BBS4, OFD1 and CEP290, are satellite clients, and their loss can disrupt satellite organization (Conkar et al., 2017; Doornbos et al., 2021; Kim et al., 2008; Kim et al., 2004; Lopes et al., 2011; Stowe et al., 2012). Whether satellite properties are altered in patient cells with ciliopathies or other disorders linked to PCM1 and its clients, and whether such phenotypes differ between clinical presentations, remains unknown.

To address this gap, we developed SatelliteQ, an open-source workflow that combines Fiji-based image processing and segmentation with Python-based data organization, spatial analysis and visualization to quantify satellite properties at the satellite and cell levels in 2D, 3D and time-lapse images. After validating its measurements against Imaris-based quantification and characterizing how segmentation settings and the choice of 2D or 3D analysis affect them, we apply SatelliteQ to cellular perturbations, different cell types, live imaging, CEP290 depletion and CEP290- and NPHP1-mutant patient-derived URECs. Together, these analyses establish SatelliteQ as an accessible framework for multidimensional satellite phenotyping across cellular contexts and disease models.

## RESULTS

### SatelliteQ is an open-source pipeline for multidimensional quantification of centriolar satellite properties

SatelliteQ is an open-source pipeline that enables quantitative measurement of centriolar satellite properties from fluorescence microscopy images through five steps (Fig. 1). It analyzes single 2D images, 3D z-stacks and 2D or 3D time-lapse series (Fig. 1, input). A satellite marker, such as PCM1 or a satellite client, is the only required channel. A cell boundary channel enables satellite measurements to be assigned to individual cells, while a centrosome channel provides a reference for centrosome-based measurements. A cilium channel can additionally be used to detect cilia and measure ciliation status and cilium length in the same cells, allowing satellite properties to be compared with cilium presence, absence and length. SatelliteQ quantifies satellite number, size (area in 2D or volume in 3D), fluorescence intensity, distance to the centrosome, shape (circularity in 2D or sphericity in 3D) and spatial distribution (Fig. 1, quantification).

**Figure 1.**
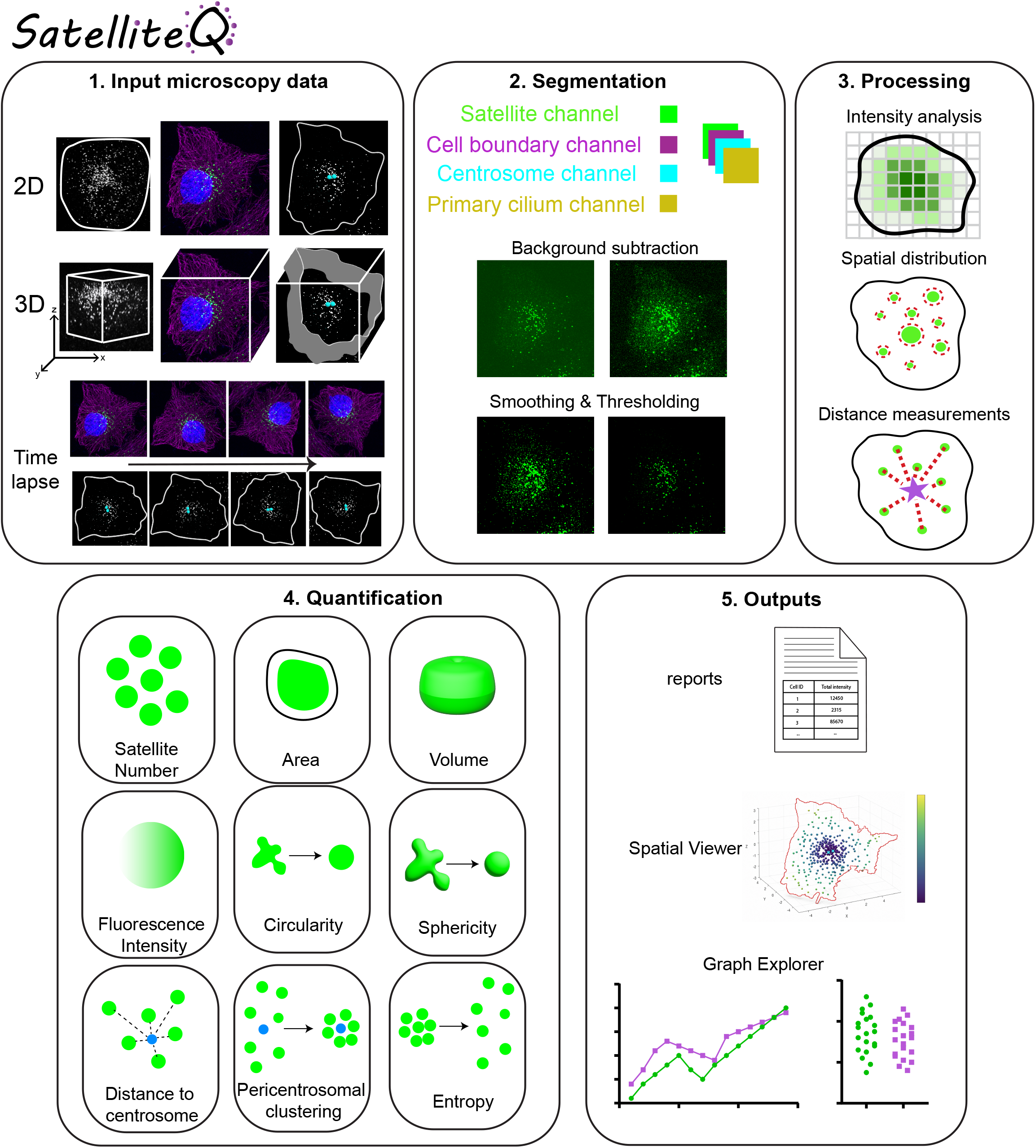
SatelliteQ workflow for quantification of centriolar satellite properties. Overview of the SatelliteQ pipeline. **(1)** Input, single 2D images, 3D z-stacks or time-lapse series containing a satellite channel and, optionally, cell boundary, centrosome and primary cilium channels. **(2)** Segmentation, each channel is processed independently through background subtraction, smoothing and thresholding to generate object masks. **(3)** Processing, segmented satellites are assigned to cells and centrosomes, and intensity, spatial distribution and distance measurements are performed. **(4)** Quantification, satellite number, area (2D), volume (3D), fluorescence intensity, distance to the centrosome (when a centrosome marker is provided), circularity (2D), sphericity (3D), spatial entropy and the pericentrosomal clustering index. When a cilium marker is provided, primary cilium count and length are also reported. **(5)** Outputs, standardized Excel reports, the interactive Spatial Viewer displaying each satellite at its measured position, and the Graph Explorer for plotting any measurement across conditions.

A Fiji macro performs image processing and segmentation (Fig. 1, segmentation and processing) (Schindelin et al., 2012), after which a Python module organizes the measurements and generates three outputs (Fig. 1, outputs). The standardized report contains satellite-, cell-, image- and, for time-lapse data, frame-level measurements. The Graph Explorer generates interactive graphs for comparing measurements across conditions and replicates, and the Spatial Viewer reconstructs individual cells and displays satellites at their measured positions. Together, these components allow SatelliteQ to quantify satellite phenotypes from individual satellites to cell- and population-level measurements.

### SatelliteQ segments satellites, cells and centrosomes and quantifies satellite properties at the satellite, cell and image levels

Quantification of satellite properties begins with the segmentation of the structures from which these measurements are derived. The Fiji macro processes the satellite, cell boundary and centrosome channels independently and then combines them (Fig. 2). The satellite channel is duplicated to preserve the original image data, background is subtracted using the rolling-ball algorithm, and the image is thresholded to generate a binary satellite mask. Thresholding uses one of the automatic methods provided by Fiji or a manual value, with the selected threshold applied consistently across all images in an experiment. Because satellite intensity and background differ between cell types, markers, treatments and microscopes, the thresholding method and the rolling-ball radius are the two parameters that most strongly affect segmentation (Fig. S4). Objects in the mask are identified with the Analyze Particles function in 2D or the 3D Objects Counter+ plugin in 3D. User-defined minimum and maximum size filters exclude objects arising from noise or aberrantly large structures, objects touching the stack edges can be excluded in 3D, and an optional watershed step separates touching satellites (Fig. 2, satellite segmentation). Each accepted object is assigned a unique Satellite ID together with its position, boundary, size and fluorescence intensity. Fluorescence intensity is measured on the background-subtracted image within the segmented object boundary and therefore reflects the fluorescence signal associated with each object. In 2D, satellites are quantified from maximum intensity projections. In 3D, they are detected across the z-stack and their 3D properties are retained. Time-lapse modes repeat the 2D or 3D analysis for each frame.

**Figure 2.**
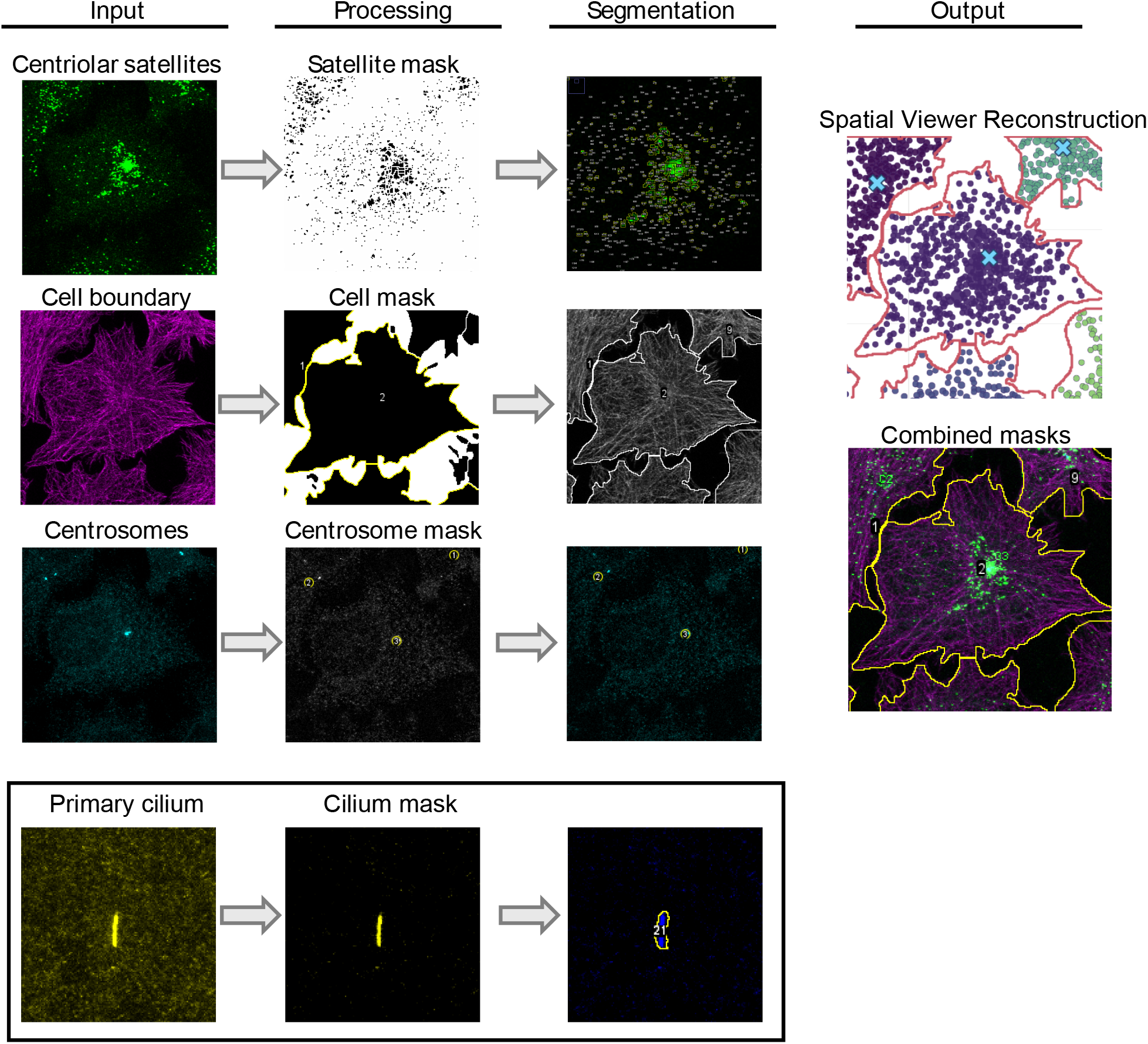
SatelliteQ segments satellites, cells and centrosomes in a single workflow. Representative fluorescence images illustrating the processing and segmentation steps of SatelliteQ. Input images of RPE1 cells contain satellites (PCM1, green), a cytoplasmic marker used for the cell boundary (α-tubulin, magenta) and centrosomes (Centrin 2, cyan). Each channel is processed independently, shown as the input, processing, segmentation and output columns. Processing generates a mask for each channel through background subtraction and thresholding, and segmentation identifies individual satellites, cell regions and centrosome positions from the masks. The output combines the three channels, assigning each satellite to a cell and a centrosome; the output images show the Spatial Viewer reconstruction and the combined masks. Primary cilium analysis is shown as an additional panel from a different representative image (ARL13B, yellow); the cilium mask identifies individual cilia. Scale bar: 10 µm.

The cell boundary is defined in one of three ways; it is segmented from a membrane or cytoplasmic marker by thresholding, approximated as a centrosome-centered circle in 2D or sphere in 3D with user-defined dimensions, or drawn manually in the ROI editor. Each cell receives a unique Cell ID. Centrosomes are detected from the centrosome channel, and their centroid coordinates serve as reference points for distance measurements. Satellites and centrosomes are then linked to the cell that contains them (Fig. 2, cell boundary and centrosome segmentation and output). Cells with more than one centrosome (e.g. after centrosome duplication or amplification) are handled by one of three user-selected rules. The user chooses whether each satellite is assigned to its nearest centrosome, whether the midpoint between the centrosomes serves as a single reference point, or whether the cell is excluded from centrosome-based measurements. Cells without a detected centrosome retain their other satellite measurements, while centrosome-based measurements are reported as missing values. Cells that are incompletely captured at the image borders are excluded during manual cell selection and ROI editing. Before quantification, all detected satellites are displayed as numbered objects overlaid on the input image. Detections that do not match the underlying fluorescence signal can be corrected manually. False positives can be deleted, merged or split objects edited, and missed satellites added. Segmentation can also be repeated with different parameters, and overlay images of the detected objects are saved for quality control. All user-defined settings are recorded with the output to support reproducibility.

SatelliteQ reports satellite properties at three levels, satellite, cell and image (Table S1). For each satellite, it records size (area in 2D or volume in 3D), integrated and mean fluorescence intensity, distance to the assigned centrosome and shape. For each cell, it reports satellite number, total and mean satellite size, total and mean satellite intensity, centrosome number, cell size and the spatial distribution measurements described in the next section. For each image, and for each frame in time-lapse data, it reports the same measurements averaged across cells. Satellite number, size and fluorescence intensity provide complementary readouts of satellite abundance and biogenesis, and their derivation from the segmented objects is illustrated in Fig. S1. Shape is reported as circularity in 2D and sphericity in 3D, which approach 1 for round objects (Fig. S1). Because most satellites are close to the resolution limit, shape measurements are most informative when satellites form larger granules, such as in cells expressing PCM1 truncation mutants (Begar et al., 2026).

The Python module reads the Fiji output, combines images from the same condition and replicate using the folder structure defined by the user, and writes a single Excel report with separate sheets for satellite-level, cell-level, image-level and, for time-lapse data, frame-level measurements. Each row is labeled with its condition, replicate, image, frame, cell and satellite identifiers. The Graph Explorer and Spatial Viewer provide interactive visualization of these data. The Graph Explorer plots measurements across conditions or replicates as distributions or means ± SD with sample size, normalizes values to a reference condition, and plots pairs of cell-level measurements with a linear fit and R². The Spatial Viewer reconstructs each cell from the segmented objects, showing satellites at their measured positions together with the cell boundary and centrosome, and can color satellites by measurements such as size, intensity or distance. Together, these outputs allow satellite properties to be examined across scales, from individual satellites within a cell to cell-level and population-level measurements.

### SatelliteQ quantifies satellite spatial organization with three complementary measurements

Satellites can cluster around the centrosome, disperse throughout the cytoplasm, or accumulate at other cellular sites, such as non-centrosomal microtubule-organizing centers or organelles to which they are experimentally repositioned (Kubo and Tsukita, 2003; Odabasi et al., 2020). Because these spatial patterns vary between cell types and change upon perturbation, no single measurement captures the full range of satellite distributions. SatelliteQ therefore describes satellite spatial organization using three complementary measurements (Fig. S2). Satellite-to-centrosome distance is the straight-line distance between the centroid of each satellite and its centrosome, measured in 2D or 3D according to the analysis mode, and the mean distance is then calculated for each cell (Fig. S2A). The pericentrosomal clustering index (PCI) also uses the centrosome position and measures the enrichment of integrated satellite fluorescence intensity around the centrosome relative to the surrounding cytoplasm, calculated within fixed radii of 3 and 12 µm (Fig. S2B). Spatial entropy is independent of the centrosome and measures how evenly satellite fluorescence is distributed throughout the cell, ranging from 0 when signal is concentrated in a single region to 1 when it is uniformly distributed (Fig. S2C). RPE1 cells spanning clustered to dispersed satellite phenotypes and the relationships among the three spatial measurements are shown in Fig. S2, D and E.

The three measurements provide distinct signatures for different types of satellite reorganization. Dispersal throughout the cytoplasm increases satellite-to-centrosome distance and entropy and lowers PCI. Tighter pericentrosomal clustering lowers distance and entropy and raises PCI. In contrast, clustering at a site away from the centrosome increases distance but lowers both entropy and PCI. Thus, PCI is particularly sensitive to pericentrosomal clustering, whereas the combination of distance and entropy helps distinguish cytoplasmic dispersal from redistribution to other cellular regions.

### SatelliteQ measurements correlate closely with those obtained by Imaris

To test whether SatelliteQ reproduces measurements obtained with established commercial software, we analyzed the same 3D images of RPE1 cells stained for PCM1, acquired for this study, with the 3D workflow of SatelliteQ and with the Imaris pipeline we used previously for object-based quantification of satellites and for distinguishing satellite phenotypes (Fig. 3A) (Begar et al., 2026). Because satellites have no ground-truth number or size, this comparison assesses agreement between the two methods rather than accuracy against a reference standard. Five properties were compared for each cell; satellite number, total satellite intensity, total satellite volume, mean satellite volume and mean distance to the centrosome. Values obtained by the two methods differed only slightly, with mean percent differences of 7.83% or less across the five properties (Fig. 3B). At the level of individual cells, satellite number, total intensity and total volume measured by SatelliteQ correlated closely with the Imaris values (R² = 0.96, 0.98 and 0.97, respectively; Fig. 3C–E). Mean satellite volume also correlated well (R² = 0.90; Fig. 3F), whereas distance to the centrosome showed greater variability but remained positively correlated (R² = 0.76; Fig. 3G). Distance is the only measurement that depends on two independently segmented objects and therefore incorporates differences in both satellite and centrosome segmentation and reference point definition. Its relatively narrow range across cells may also contribute to the lower R². When measurements from all cells were plotted side by side, SatelliteQ closely reproduced the cell-to-cell variation measured by Imaris for all five properties (Fig. S3A–E). Thus, SatelliteQ reproduces Imaris measurements across satellite properties while providing an automated and freely available alternative for object-based satellite quantification.

**Figure 3.**
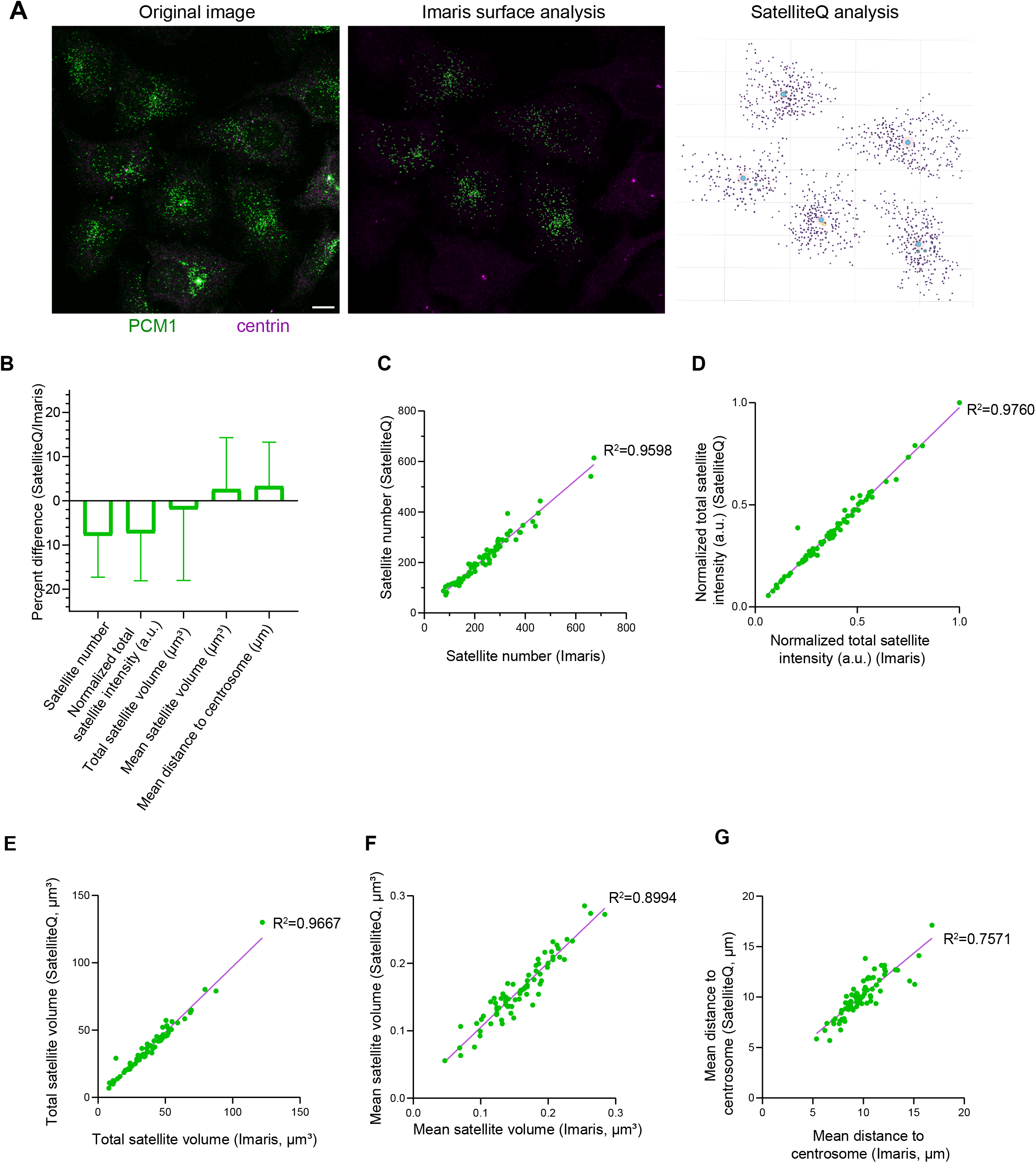
SatelliteQ measurements correlate closely with those obtained by Imaris. **(A)** Overview of the comparison. The same 3D images of RPE1 cells stained for PCM1 (green) and Centrin2 (magenta) were analyzed independently in 3D with Imaris (surface analysis) and SatelliteQ (Spatial Viewer reconstruction) using matched settings (Table S2). **(B)** Percent difference between SatelliteQ and Imaris values per cell for the five properties compared. Bars represent mean ± SD across cells. **(C to G)** Values from SatelliteQ plotted against values from Imaris for each cell, for satellite number **(C)**, total satellite fluorescence intensity **(D)**, total satellite volume **(E)**, mean satellite volume **(F)** and mean satellite-to-centrosome distance **(G)**. Lines show linear fits and R² values are indicated. Each dot represents one cell. Intensity values are normalized to the maximum value of each replicate, set to 1. Results are from 2 biological replicates (73 cells in total). Scale bar: 10 µm.

### Segmentation parameters and the choice of 2D or 3D analysis affect satellite measurements

There is not an established ground truth measurement for satellite number or size in fluorescence microscopy. Individual satellites are reported to be approximately 70 to 100 nm electron-dense granules by electron microscopy (Kubo et al., 1999). These structures are below the resolution limit of conventional light microscopy, so fluorescence images report apparent rather than ultrastructural satellite sizes. The number of detected satellites likewise depends on how individual objects are segmented. We therefore asked how strongly segmentation settings and the choice of 2D or 3D analysis influence SatelliteQ measurements. To address this, we analyzed six images of RPE1 cells, each containing multiple cells, in 2D and 3D, varying one segmentation parameter at a time (Fig. S4).

In 2D, the thresholding method strongly affected satellite number and total area (Fig. S4A and B). Lower manual threshold values detected more satellites and a larger total satellite area, with the additional objects predominantly small (Fig. S4D). Among the automatic methods, Triangle and Li detected more objects than Otsu, Moments, Yen and the hysteresis-like method, and the size distributions again showed that these additional objects were predominantly small (Fig. S4C). In contrast, changing the rolling-ball radius had little effect on satellite number, total area or the size distribution (Fig. S4E–G). Watershed separation produced only modest changes, slightly increasing satellite number and decreasing total area, consistent with the splitting of touching objects (Fig. S4H and I). In 3D, satellite number and total satellite volume decreased as the threshold increased (Fig. S4J and K), with the additional objects at low thresholds again predominantly small (Fig. S4L), whereas changing the rolling-ball radius had relatively little effect (Fig. S4M–O). Thus, thresholding was the dominant source of variation among the parameters tested, primarily by changing the detection of small, low-intensity objects, whereas background subtraction and watershed separation had comparatively limited effects within the tested ranges.

We next compared the 2D and 3D analysis modes directly, using a separate set of images of RPE1 cells stained for PCM1 to mark satellites and α-tubulin to mark microtubules, with each image analyzed in both modes (Fig. 4A). Satellite number per cell was similar in the two modes, with a trend toward higher counts in 3D (Fig. 4B), consistent with Fig. S4, in which 3D analysis detected more satellites than 2D across the tested settings. Total satellite size is reported as area in 2D and as volume in 3D and is therefore interpreted separately within each mode (Fig. 4C and D).

**Figure 4.**
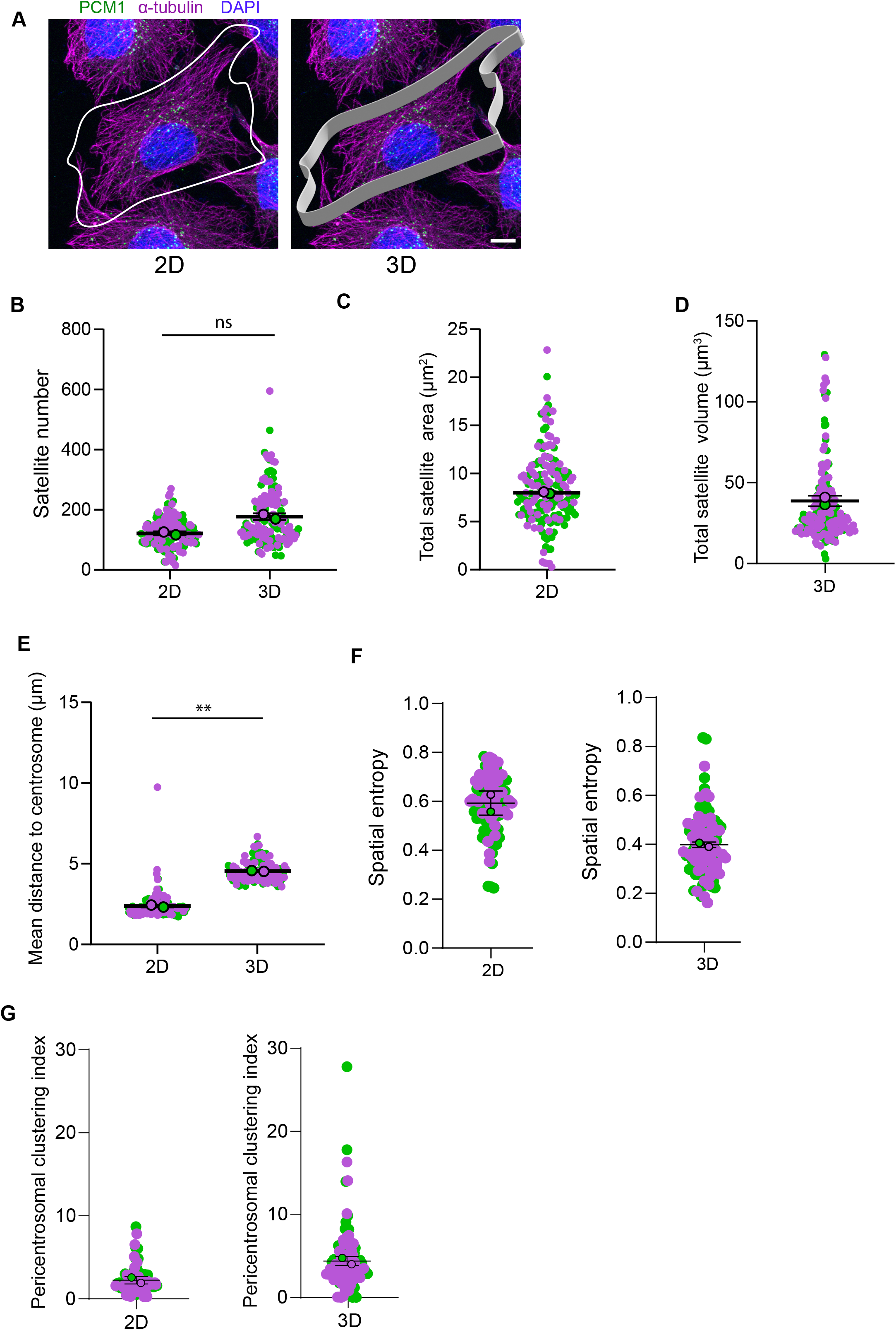
Comparison of satellite measurements in the 2D and 3D analysis modes of SatelliteQ. **(A)** Representative image of RPE1 cells stained for PCM1 (green), α-tubulin (magenta) and DAPI (blue), analyzed with the 2D mode using a manually defined cell boundary (white outline) and with the 3D mode using a rendered cell boundary (grey band). **(B)** Satellite number per cell in the two modes. **(C and D)** Total satellite area per cell in 2D **(C)** and total satellite volume per cell in 3D **(D)**; size is interpreted within each mode. **(E to G)** Mean satellite-to-centrosome distance **(E)**, spatial entropy **(F)** and pericentrosomal clustering index **(G)** per cell in the two modes. The same cells were analyzed in both modes. Each dot represents one cell; colours indicate biological replicates and larger outlined circles represent replicate means. Black bars represent mean ± SD. Results are from 2 biological replicates (more than 72 cells per replicate). Statistical significance was determined by paired two-tailed Student’s t test; *, P < 0.05; ***, P < 0.001; ns, not significant. Scale bar: 10 µm. Analysis settings are listed in Table S2.

The two modes gave systematically different values for the three spatial organization measurements. Distances measured in 2D were substantially smaller than in 3D (Fig. 4E). This difference likely reflects the loss of the z component in projected distances; satellites that appear close to the centrosome in a projection can be farther away when their full xyz coordinates are considered. Spatial entropy was higher and the pericentrosomal clustering index lower in 2D than in 3D (Fig. 4F and G). These differences are consistent with the loss of three-dimensional spatial information when the z dimension is collapsed, which alters how satellite signal is distributed across the analyzed cell region. The absolute values of the spatial measurements are therefore mode-dependent and should be compared only within the same analysis mode.

Together, these results show that segmentation settings influence the absolute values of satellite measurements, while 3D analysis provides spatial information that is lost in 2D projections. Absolute values obtained with different settings should therefore not be treated as interchangeable, and 2D and 3D size measurements should not be compared directly because they report area and volume, respectively. For comparisons between experimental conditions, segmentation settings should be optimized for each marker, cell type and imaging setup and then kept consistent across the dataset. The same analysis mode should likewise be used across the conditions being compared. The segmentation settings used for each analysis in this study are listed in Table S2.

### SatelliteQ captures distinct changes in satellite properties induced by cellular perturbations

We next asked whether SatelliteQ could resolve biologically distinct changes in satellite properties following perturbations known to affect satellite abundance and spatial organization. RPE1 cells were treated with DMSO as control, with the microtubule-depolymerizing drug nocodazole or with 1,6-hexanediol (1,6-HD), which disrupts weak hydrophobic interactions, and PCM1-positive satellites were quantified in 3D (Fig. 5A). Both treatments reduced satellite number, total satellite volume and total satellite intensity compared with control cells (Fig. 5B–D). These reductions are consistent with our previous identification of a 1,6-HD-sensitive satellite population and with the requirement for an intact microtubule network for the movement, pericentrosomal clustering and biogenesis of satellites (Begar et al., 2026; Conkar et al., 2019; Dammermann and Merdes, 2002; Kubo et al., 1999). Thus, both perturbations reduced satellite abundance under the conditions tested.

**Figure 5.**
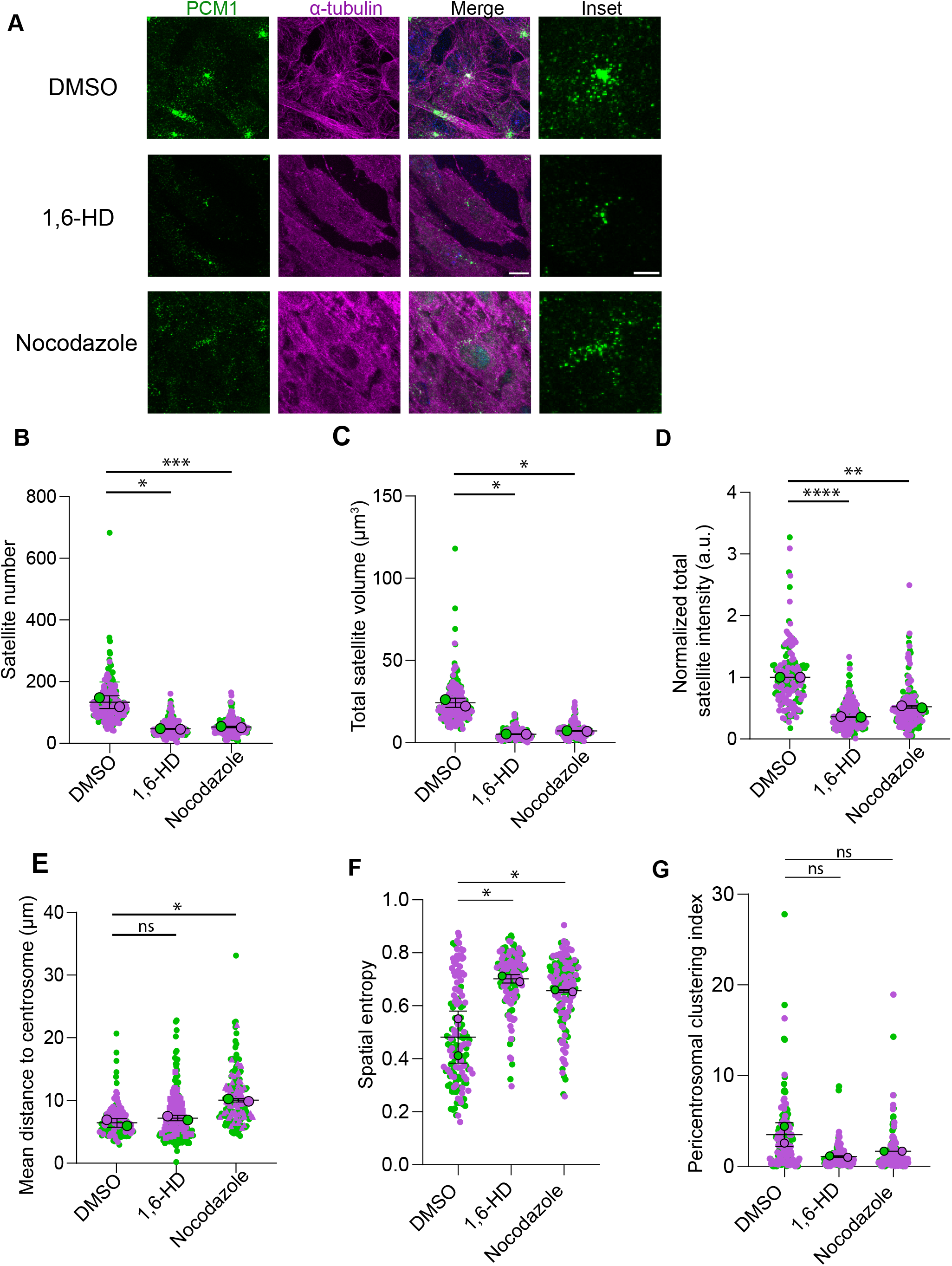
SatelliteQ captures changes in satellite properties induced by nocodazole and 1,6-hexanediol. **(A)** Representative images of RPE1 cells treated with DMSO, 5% 1,6-hexanediol (1,6- HD) for 5 min or 5 µg/ml nocodazole for 1 h, stained for PCM1 (green), α-tubulin (magenta) and DAPI (blue). Insets show enlarged views of the pericentrosomal region. **(B to G)** Satellite number **(B)**, total satellite volume **(C)**, normalized total satellite fluorescence intensity **(D)**, mean satellite-to-centrosome distance **(E)**, spatial entropy **(F)** and pericentrosomal clustering index **(G)** per cell for each treatment. Intensity values are normalized to the mean of the DMSO condition, set to 1, to allow comparison across experiments. Each dot represents one cell; colours indicate biological replicates (green and magenta) and larger outlined circles represent replicate means. Black bars represent mean ± SD. Results are from 2 biological replicates (more than 83 cells per condition per replicate). Statistical significance was determined by two-tailed unpaired Student’s t test; **, P < 0.01; ns, not significant. Scale bars: 10 µm; insets, 2 µm. Analysis settings are listed in Table S2.

We next examined the spatial organization of the remaining satellites. Nocodazole increased the mean satellite-to-centrosome distance, whereas 1,6-HD did not (Fig. 5E). Spatial entropy increased after both treatments (Fig. 5F), while the pericentrosomal clustering index showed a downward trend that did not reach statistical significance (Fig. 5G), consistent with reduced pericentrosomal enrichment. Because entropy reports the overall uniformity of satellite distribution, whereas mean distance reports radial position relative to the centrosome, these results indicate that both treatments produced a more spatially dispersed satellite distribution, but only nocodazole increased the distance of satellites from the centrosome. Thus, similar reductions in satellite abundance were accompanied by distinct changes in spatial organization that could not be distinguished by satellite number, size or intensity alone.

Because segmentation settings determine the absolute values of satellite measurements (Fig. S4), a biological difference detected with one setting could in principle be an artifact of that choice. We therefore used the nocodazole-induced satellite phenotype established above as a test case and asked whether it remains detectable across segmentation settings. Control and nocodazole-treated cells were analyzed using different thresholding methods, with each setting applied identically to both conditions (Fig. S5). The nocodazole-induced increase in mean satellite-to-centrosome distance was detected with every thresholding method in both 2D and 3D (Fig. S5A and C). Satellite number decreased after nocodazole treatment under every tested setting (Fig. S5B and D). The decrease did not reach statistical significance with some stringent thresholds, which detected very few satellites in both control and nocodazole-treated cells, or with the Triangle method in 2D. Importantly, the direction of both biological effects was preserved across all tested settings.

Together, these results show that SatelliteQ detects biologically meaningful changes in satellite number, size, fluorescence intensity and spatial organization and resolves differences in the spatial response to distinct cellular perturbations. The robustness of the nocodazole phenotype across segmentation settings further supports the use of matched segmentation parameters for quantitative comparisons between conditions.

### Satellite properties differ between cell types and satellite number scales with cell size

We next asked whether satellite properties differ between cell types. HeLa (human cervical adenocarcinoma), HEK293T (human embryonic kidney), RPE1 (human retinal pigment epithelial) and U2OS (human osteosarcoma) cells were stained for PCM1 to mark satellites, α-tubulin to mark microtubules and centrin to mark centrioles, and analyzed with the 3D workflow (Fig. 6A–D).

**Figure 6.**
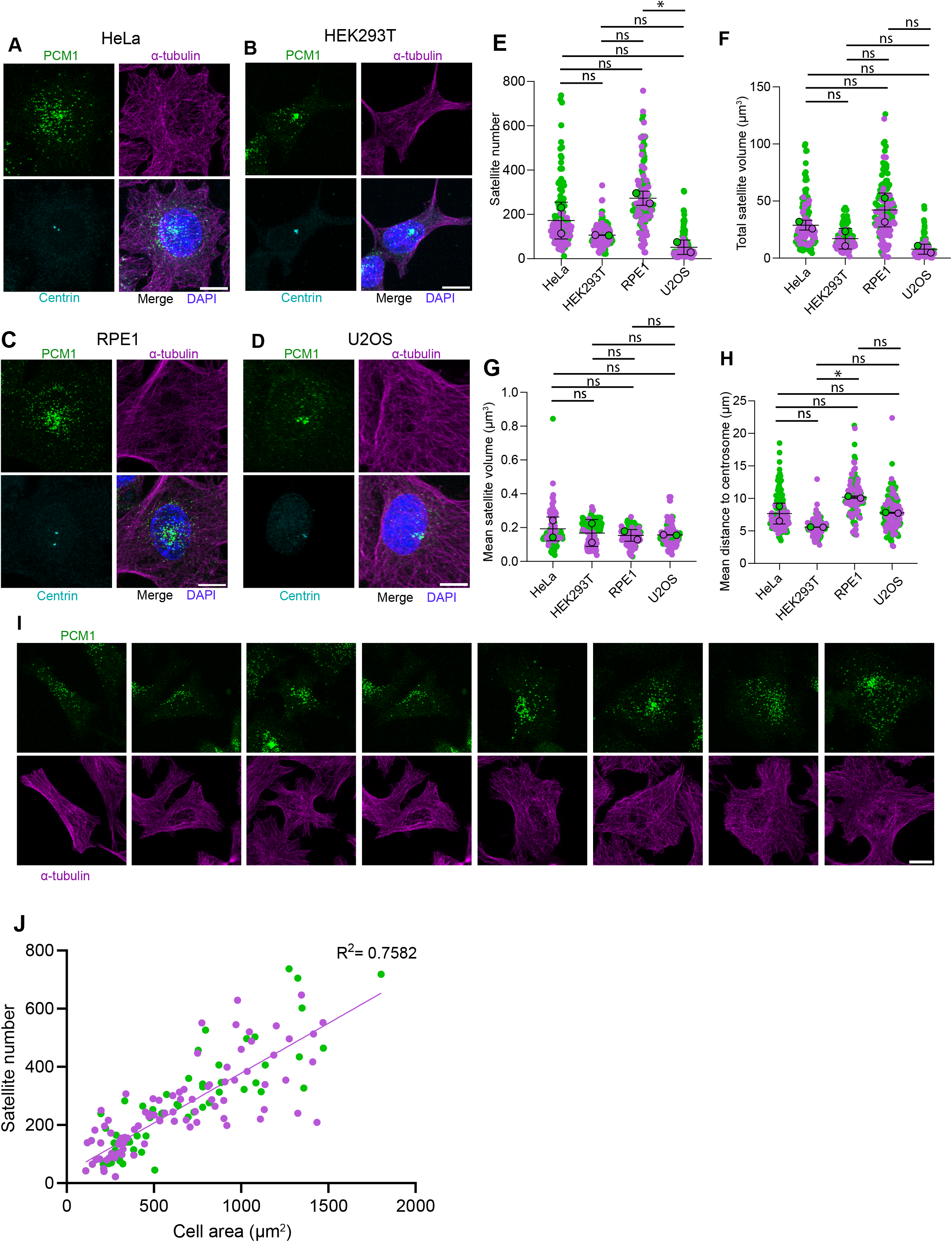
Satellite properties differ between cell types and satellite number scales with cell size. **(A to D)** Representative images of HeLa **(A)**, HEK293T **(B)**, RPE1 **(C)** and U2OS **(D)** cells stained for PCM1 (green), α-tubulin (magenta), centrin (cyan) and DAPI (blue). **(E to H)** Satellite number **(E)**, total satellite volume **(F)**, mean satellite volume **(G)** and mean satellite-to-centrosome distance **(H)** per cell, quantified with the 3D workflow. **(I)** Representative RPE1 cells of different sizes stained for PCM1 (green) and α-tubulin (magenta). **(J)** Satellite number plotted against cell area for individual RPE1 cells, quantified with the 2D workflow. The line shows a linear fit and R² is indicated. Each dot represents one cell; colours indicate biological replicates (green and magenta). Black bars represent mean ± SD. Results are from 2 biological replicates (more than 47 cells per line per replicate). Statistical significance was determined by one-way Anova test.; *, P < 0.05; **, P < 0.01; ***, P < 0.001 ; ns, not significant. Scale bars: 10 µm. Analysis settings are listed in Table S2.

Satellite number and total satellite volume differed between the four lines; RPE1 cells contained more satellites and a larger total satellite volume than HEK293T and U2OS cells, with HeLa cells intermediate (Fig. 6E and F). In contrast, mean satellite volume was similar across all four lines (Fig. 6G), indicating that the differences in total satellite volume primarily reflect differences in satellite number rather than mean satellite size. Mean satellite-to-centrosome distance also varied between lines and was smallest in HEK293T cells and largest in RPE1 cells (Fig. 6H). These differences may reflect variation in cell size and shape as well as in the extent of pericentrosomal satellite clustering between lines. Cell size alone did not explain the differences across cell lines, as U2OS cells are larger and flatter than HEK293T cells yet contained fewer satellites.

However, satellite number varied substantially between individual cells within each cell line (Fig. 6E). We therefore asked whether cell size could account for this heterogeneity. In RPE1 cells spanning a wide range of sizes (Fig. 6I), satellite number scaled with cell area (R² = 0.76; Fig. 6J), indicating that cell size is a major correlate of satellite number within this population. Together, these results show that SatelliteQ resolves differences in satellite properties between cell types and reveals cell-size-dependent variation in satellite number within a cell population.

### SatelliteQ quantifies the formation, remodeling and morphology of PCM1 granules

We next asked whether SatelliteQ quantifies dynamic changes in satellites during live-cell imaging. Because each frame of a time-lapse series is segmented and quantified independently, changes in satellite properties can be followed over time without tracking individual objects between frames. We first analyzed time-lapse images of RPE1 PCM1 knockout cells expressing doxycycline-inducible GFP-PCM1-N, an N-terminal fragment of PCM1 (amino acids 1 to 1200), which forms larger granules than the full-length protein and is partially functional in ciliogenesis assays (Begar et al., 2026; Wang et al., 2016). Because its larger granules are easier to resolve, PCM1-N has been used in parallel with full-length PCM1 to study granule biogenesis, and its induction allows de novo granule formation to be followed from the start (Begar et al., 2026). After induction, GFP-PCM1- N granules appeared and rapidly increased in number, and granule number then gradually declined while mean granule area and total fluorescence intensity continued to increase (Fig. 7A–D). These kinetics reproduce the biogenesis phases we defined previously, in which an initial nucleation phase is followed by growth and by fusion of granules that reduces their number over time.

**Figure 7.**
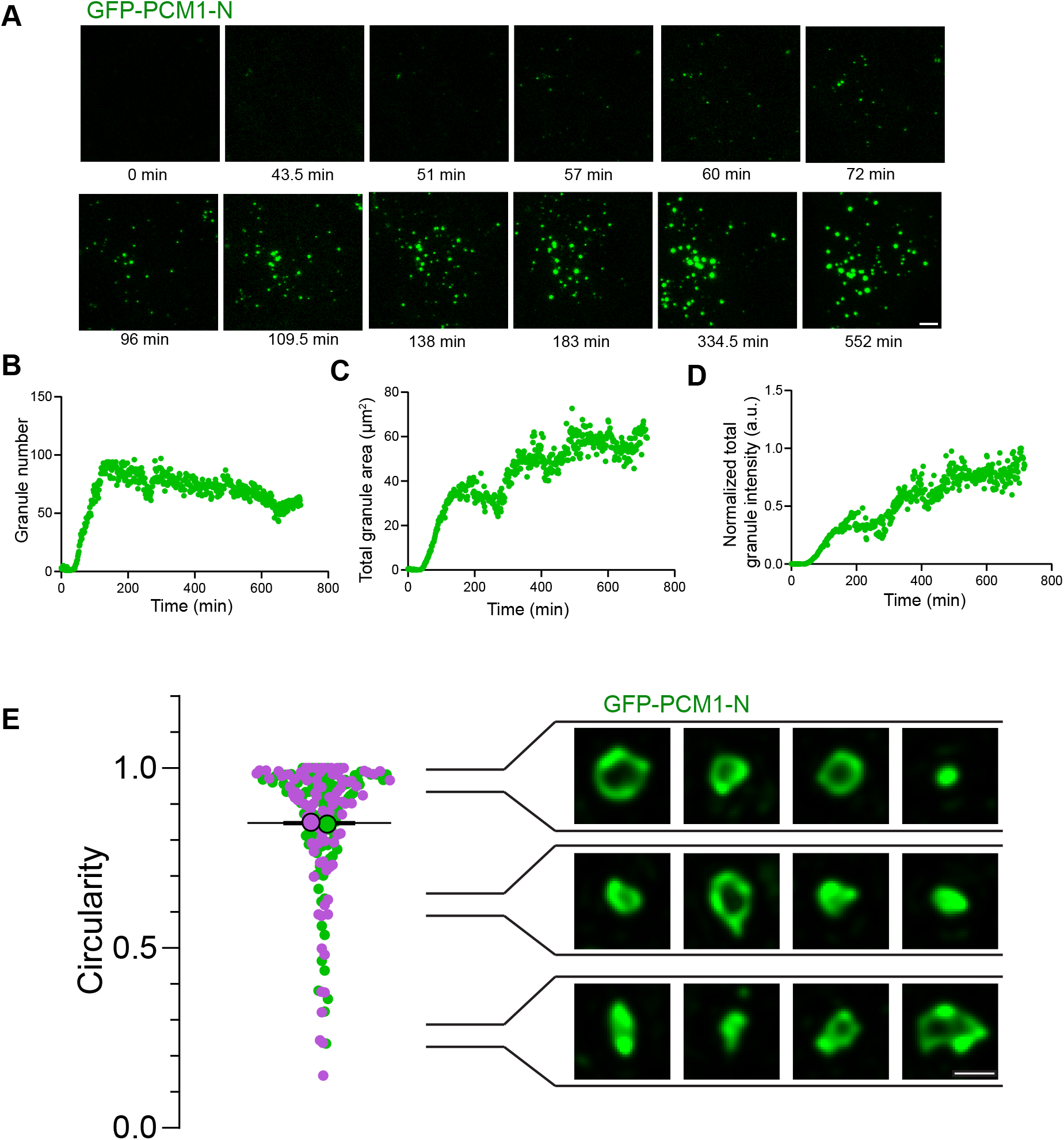
SatelliteQ quantifies the formation, remodeling and shape of PCM1 granules in live cells. **(A)** Stills from time-lapse imaging of an RPE1 PCM1 knockout cell expressing doxycycline-inducible GFP-PCM1-N; times are relative to doxycycline addition and are indicated below each still. **(B to D)** Granule number **(B)**, total granule area **(C)** and normalized total granule fluorescence intensity **(D)** over time for the same representative cell. **(E)** Circularity of GFP-PCM1-N granules imaged by structured illumination microscopy. Insets show representative granules at the indicated circularity ranges, from round to elongated and irregular. Each dot represents one granule; colours indicate biological replicates (green and magenta). Black bars represent mean ± SD. Results are from 2 biological replicates (more than 44 granules per replicate). Intensity values are normalized to the maximum fluorescence intensity set to 1. Scale bars: 10 µm (A); 0.5 µm (E, insets). Analysis settings are listed in Table S2.

Satellite disassembly at mitotic entry and reassembly upon mitotic exit are physiological features of satellite remodeling during cell division, and GFP-PCM1-N granules behave similarly (Begar et al., 2026; Dammermann and Merdes, 2002; Kubo and Tsukita, 2003). In cells that entered mitosis during imaging, SatelliteQ captured this dissolution as coordinated decreases in granule number, total area and total intensity, the properties that report granule abundance, followed by recovery of these measurements after cell division (Fig. S6A–D). Because each frame is quantified independently and each cell is analyzed separately, these dissolution and reassembly cycles are resolved at the level of individual cells.

Because GFP-PCM1-N forms granules that are larger than endogenous satellites, they also allow shape quantification. We analyzed structured illumination microscopy images of GFP-PCM1-N granules and examined their circularity, which approaches 1 for circular objects and decreases for irregular shapes (Fig. 7E). Circularity values closely reflected the appearance of individual granules, with near-circular outlines giving high values and elongated or irregular granules giving lower values across the measured range. Together, these results show that SatelliteQ quantifies the formation, remodeling and morphology of granules in living cells.

### CEP290 depletion alters satellite abundance and spatial organization in RPE1 cells

CEP290 is a satellite client that is recruited to the PCM1 scaffold after its formation during satellite biogenesis, and its localization within satellites is remodeled during cilium formation (Begar et al., 2026). CEP290 mutations cause a broad spectrum of ciliopathies ranging from isolated retinal degeneration to Joubert syndrome and nephronophthisis (Coppieters et al., 2010). We therefore asked whether perturbing a client alters satellite properties, using CEP290 as a disease-relevant example. CEP290 was depleted with a validated siRNA, and depletion was confirmed by immunofluorescence (Fig. S7A). Its depletion also reduced ciliation after serum starvation, consistent with its established role in ciliogenesis (Fig. S7B) (Kim et al., 2008). SatelliteQ analysis of PCM1-stained cells in 3D revealed changes in several satellite properties (Fig. 8A). Satellite number, total PCM1 intensity and total satellite volume were reduced, indicating reduced satellite abundance (Fig. 8B–D). The pericentrosomal clustering index increased upon CEP290 depletion, whereas mean satellite-to-centrosome distance and spatial entropy decreased slightly (Fig. 8E–G). Together, these three measurements indicate increased pericentrosomal clustering of the remaining satellites, consistent with previous reports (Firat-Karalar, 2018; Kim et al., 2008). The spatial phenotype is the converse of the dispersal observed after nocodazole treatment, for which distance and entropy provided the more sensitive measurements (Fig. 5E–G). Thus, although CEP290 is recruited to the PCM1 scaffold as a client, its loss alters both the abundance and spatial organization of satellites, indicating that satellite properties depend on satellite clients in addition to the PCM1 scaffold itself.

**Figure 8.**
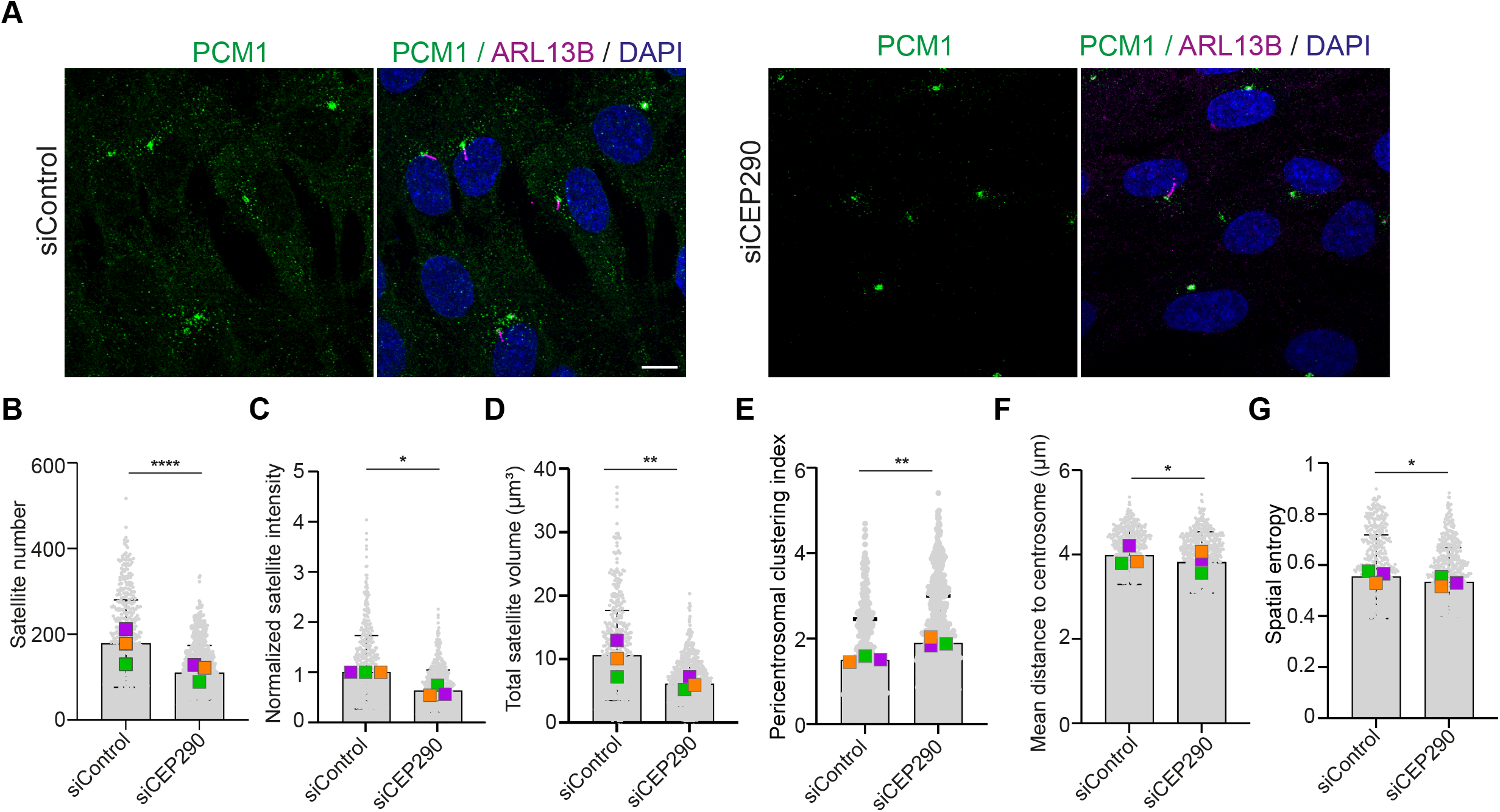
CEP290 depletion alters satellite abundance and spatial organization in RPE1 cells. **(A)** Representative images of RPE1 cells transfected with control or CEP290 siRNA, serum starved for 24 h, and stained for PCM1 (green), ARL13B (magenta) and DAPI (blue). **(B to G)** Satellite number **(B)**, normalized total satellite fluorescence intensity **(C)** total satellite volume **(D)**, pericentrosomal clustering index **(E)**, mean satellite-to- centrosome distance **(F)** and spatial entropy **(G)** per cell, quantified with the 3D workflow. Intensity values are normalized to the mean of the siControl condition, set to 1. Each dot represents one cell; coloured squares represent replicate means. Bars represent mean ± SD. Results are from 3 biological replicates (minimum 120 cells per condition). Statistical significance was determined by unpaired two-tailed Student’s t test; *, P < 0.05; **, P < 0.01; ****, P < 0.0001. Scale bar: 10 µm. Analysis settings are listed in Table S2.

Because satellites are remodeled during cilium formation and CEP290 depletion also reduces ciliation, we next asked how satellite properties relate to ciliation status at the single cell level. Serum-starved wild-type RPE1 cells stained for PCM1, ARL13B and γ- tubulin were classified as ciliated or unciliated with the cilium detection module, and the satellite properties of the two groups were compared in the same images (Fig. S7C). Cells without cilia contained more satellites with higher normalized intensity but similar total satellite volume, together with a lower pericentrosomal clustering index, a larger mean distance to the centrosome and higher spatial entropy (Fig. S7D–I). Thus, unciliated cells carry more abundant and more dispersed satellites, whereas ciliated cells concentrate fewer satellites around the centrosome. Notably, this is the opposite of the CEP290 depletion phenotype; if the satellite changes in CEP290-depleted cells were a secondary consequence of reduced ciliation, they should have shifted toward the more abundant, more dispersed pattern of unciliated cells. The satellite phenotype of CEP290-depleted cells therefore does not simply reflect their ciliation state.

### Satellite properties are altered in CEP290-mutant patient-derived URECs

CEP290 depletion showed that the loss of a single client is sufficient to disrupt satellite homeostasis (Fig. 8). Because many satellite clients, unlike PCM1 itself, are mutated in ciliopathies, satellite properties may provide an additional organellar phenotype for assessing defects of the centrosome-cilium-satellite axis. To test this, we asked whether satellite properties are altered in patient-derived cells. We compared urine-derived renal epithelial cells (URECs) from two healthy donors, two patients carrying CEP290 mutations, one with Leber congenital amaurosis (LCA) with retinal and renal involvement (CEP290 mutant 1) and one with Senior–Løken syndrome (SLS) with renal involvement (CEP290 mutant 2), and two patients carrying NPHP1 mutations, both with renal involvement (nephronophthisis) (Garcia et al., 2022; Malglaive et al., 2025; Menguy et al., 2026). Cells were stained for PCM1, γ-tubulin to mark centrosomes and acetylated tubulin to mark cilia and analyzed with the 2D workflow (Fig. 9A). All UREC lines were analyzed in the same mode with identical settings.

**Figure 9.**
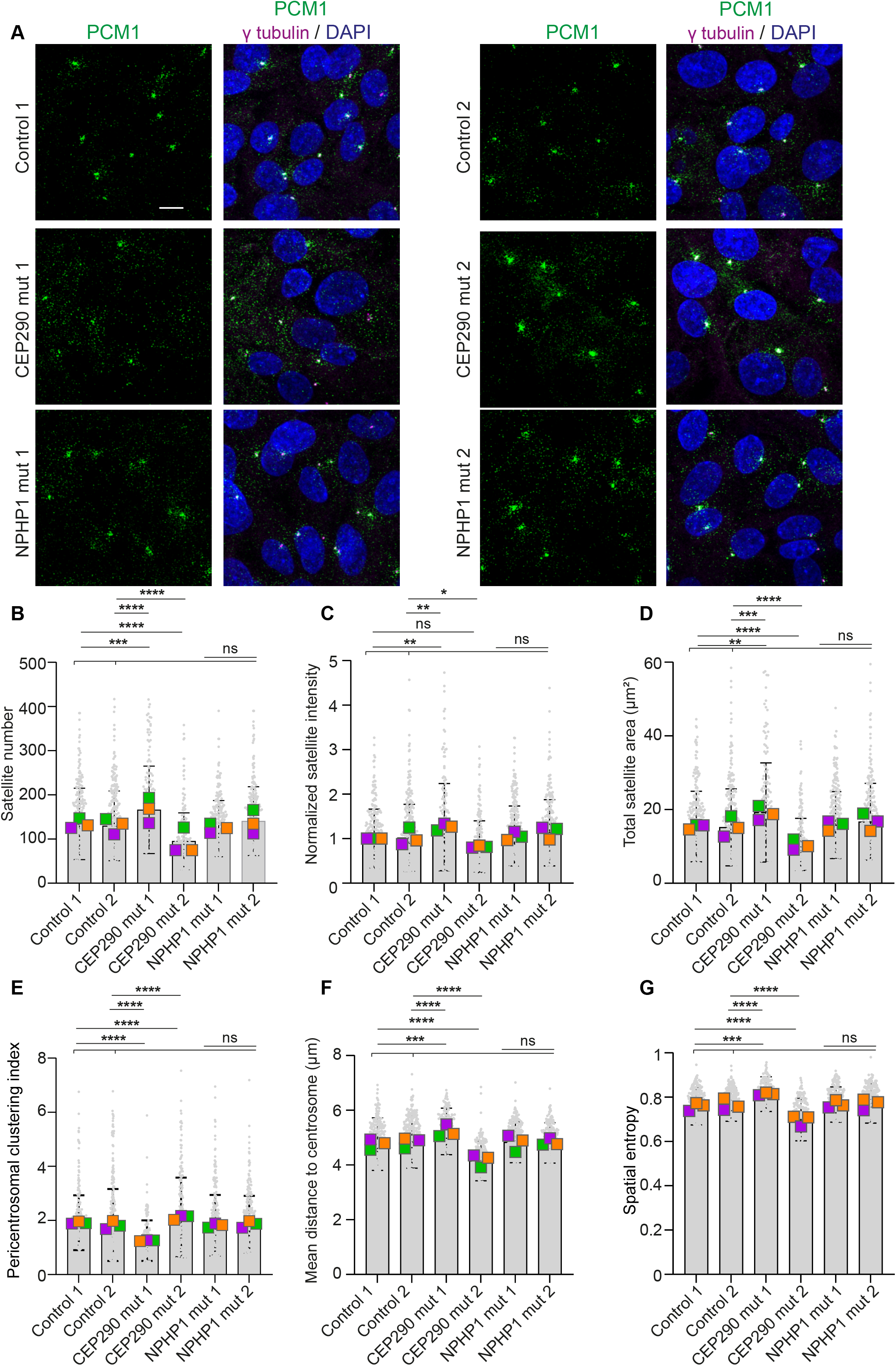
Satellite properties are altered in CEP290-mutant patient-derived URECs. **(A)** Representative images of URECs from two healthy donors (Control 1 and 2), two patients with CEP290 mutations and two patients with NPHP1 mutations, stained for PCM1 (green), γ-tubulin (magenta) and DAPI (blue). **(B to G)** Mean satellite-to-centrosome distance **(B)**, spatial entropy **(C)**, satellite number **(D)**, normalized total satellite fluorescence intensity **(E)**, total satellite area **(F)** and pericentrosomal clustering index **(G)** per cell, quantified with the 2D workflow. Intensity values are normalized to the mean of Control 1, set to 1. Each dot represents one cell; coloured squares represent replicate means. Bars represent mean ± SD. Brackets show comparisons of each CEP290-mutant line with each control line; ns spans the comparisons of the NPHP1- mutant lines with the controls. Results are from 3 biological replicates (minumum 50 cells per line). Statistical significance was determined by one-way ANOVA with multiple comparisons; *, P < 0.05; **, P < 0.01; ***, P < 0.001; ****, P < 0.0001; ns, not significant. Scale bar: 10 µm. Analysis settings are listed in Table S2.

Strikingly, both CEP290-mutant lines showed altered satellite properties compared with the controls, but in opposite directions. CEP290 mutant 1 showed increased satellite number, total intensity, total area, centrosome distance and spatial entropy, together with a reduced pericentrosomal clustering index, indicating more abundant and more dispersed satellites (Fig. 9B–G). In CEP290 mutant 2, these changes were reversed, indicating fewer satellites with lower total intensity and area and increased pericentrosomal clustering (Fig. 9B–G). The divergent satellite phenotypes may reflect the different CEP290 variants or genetic backgrounds of the two patients. The NPHP1- mutant lines did not differ significantly from the controls across the measured satellite properties (Fig. 9B–G), indicating that altered satellite properties are not a uniform feature of the ciliopathy UREC lines examined here.

We next asked whether the divergent satellite phenotypes correlate with ciliation, the classical functional readout of ciliopathy cells. Ciliation was reduced in both CEP290- mutant lines and in NPHP1 mutant 2, whereas NPHP1 mutant 1 ciliated at control levels (Fig. S8A, S8B). Satellite phenotypes did not simply mirror these defects; the same reduction in ciliation accompanied opposite satellite phenotypes in the two CEP290 lines and no significant satellite changes in NPHP1 mutant 2. Satellite properties therefore capture information beyond ciliation alone, adding a dimension to quantifying defects of the centrosome-cilium-satellite axis. Together, these results provide quantitative analysis of satellite properties in patient-derived cells and show that satellite phenotypes can arise both after acute depletion of a satellite client and in cells carrying disease-associated mutations.

## Discussion

SatelliteQ brings multidimensional, object-based quantification of centriolar satellites, previously restricted to commercial software and custom pipelines, into an open-source workflow. It measures every satellite as an individual object in 2D, 3D and time-lapse images and reports abundance, size, intensity, shape and spatial organization at the satellite, cell and image levels. This resolution detected phenotypes that single measurements would have missed, from treatment-specific dispersal signatures to divergent satellite phenotypes in patient-derived cells carrying CEP290 mutations. Absolute values depend on segmentation settings, but the biological differences tested here were preserved when matched settings were applied across conditions. Built on Fiji and Python with a graphical interface and interactive outputs, SatelliteQ requires no programming and no commercial license.

Quantitative image analysis tools have transformed the study of organelles with well-defined morphological features. For cilia, CiliaQ, ACDC and detectCilia standardize morphological and intensity measurements across laboratories, and automated cilium phenotyping supports ciliopathy diagnosis in patient cells (Budde-Sagert et al., 2024; Doornbos et al., 2021; Hansen et al., 2021; Lauring et al., 2019). For centrioles and centrosomes, automated tools count centrioles and quantify centrosomes together with microtubule organization (Burgy et al., 2023; Sankaran et al., 2020). Single-cell centrosome quantification has also been established in normal and cancer tissue (Wang et al., 2019). No comparable tool has existed for satellites, whose diversity has instead been collapsed into qualitative categories or a single pericentrosomal intensity value. As membrane-less organelles emerge as a central theme in cell biology, and as satellite dysfunction is increasingly linked to ciliopathies, neurodevelopmental disorders and cancer, standardized satellite measurement is timely. SatelliteQ extends object-based organelle phenotyping to satellites and, like the cilia tools before it, aims to make satellite phenotypes comparable across studies through recorded settings and open code.

SatelliteQ also makes satellite heterogeneity itself measurable. Satellite number, size, intensity and distribution varied between cell types, between cells of one type and within single cells over time. Correlating these measurements identified cell size as one source of this variation; satellite number scaled with cell area within a cell line, whereas mean apparent satellite size stayed similar across cell lines. Satellite number should therefore be interpreted in the context of cell size, which SatelliteQ measures in the same cells. Future studies can apply this approach by quantifying satellite properties together with cell size across conditions, cell types and disease states.

The sources of this heterogeneity remain largely unexplained. This connects satellite biology to a broader theme in cell biology; organelle heterogeneity is increasingly recognized as biological variation rather than measurement noise (Chang and Marshall, 2017). Mitochondria are morphologically and functionally heterogeneous within single cells, lysosomes form subpopulations with distinct positioning and dynamics, and automated pipelines now resolve this diversity continuously within and between individual cells (Ba et al., 2018; Bayati et al., 2026; Collins et al., 2002). SatelliteQ allows this concept of organelle heterogeneity to be examined in a membrane-less organelle. Because SatelliteQ measures every satellite as an object and every cell separately, heterogeneity between cells and between individual satellites can be quantified directly, providing a way to test how cell cycle state, differentiation and other cellular states contribute to satellite variation, as shown here for ciliation state. Satellites also differ in composition (Firat-Karalar et al., 2014; Gheiratmand et al., 2019), and combining SatelliteQ with multiplex imaging of satellite clients could extend the analysis from satellite properties to compositional heterogeneity within and between cells.

An important outcome of this study is that the three spatial measurements capture different properties of satellite organization and are complementary rather than redundant. Nocodazole, 1,6-hexanediol and CEP290 depletion each altered a different combination of centrosome distance, spatial entropy and pericentrosomal clustering. Which measurement was most informative depended on the phenotype; entropy and distance were most sensitive to dispersal, whereas pericentrosomal clustering index best captured clustering around the centrosome. Together, the three measurements distinguish perturbations that similar changes in abundance would leave indistinguishable. Thus, in our datasets, quantifying all three measures together captured phenotypes that no single measure resolved on its own.

Our results establish satellite properties as a disease-relevant readout and provide the first object-based, multidimensional quantification of satellite properties in patient-derived cells. Acute CEP290 depletion quantitatively confirmed the pericentrosomal aggregation of PCM1 granules described previously (Kim et al., 2008). In URECs, the two CEP290- mutant lines showed divergent satellite phenotypes and the NPHP1-mutant lines did not differ from the controls. Satellite phenotypes did not simply mirror changes in ciliation, suggesting that they provide information beyond ciliation alone. Ciliopathies are particularly relevant because many ciliopathy proteins are satellite clients and the same gene can cause distinct renal, retinal or syndromic presentations, as in our two CEP290 patients. Quantitative cellular assays are increasingly used to support patient classification and variant interpretation (Doornbos et al., 2021). Because SatelliteQ records cilium presence and length in the same cells, satellites can be phenotyped together with cilia, as shown by the distinct satellite organization of ciliated and unciliated RPE1 cells, a step toward quantitative readouts across the centrosome-cilium-satellite axis. Our patient data are a proof of principle; with two lines per genotype, the divergent CEP290 phenotypes motivate rather than establish satellite phenotyping as a classifier. Larger panels and accessible cells such as URECs will be needed to determine whether satellite signatures relate to genotype or clinical presentation. The relevance of satellite phenotyping may extend beyond ciliopathies. In trisomy 21, increased levels of the satellite client pericentrin cause pericentrosomal crowding, disrupt satellite localization and delay ciliogenesis (Galati et al., 2018; Jewett et al., 2023; McCurdy et al., 2022). More broadly, satellite properties may provide a quantitative readout in other disease states that perturb the centrosome-cilium-satellite axis. In disease states that alter cell size, satellite phenotypes should also be interpreted in the context of the scaling described above.

The main limitations of satellite quantification stem from the small size of satellites and their dense pericentrosomal organization. Most satellites lie near or below the diffraction limit, so the sizes reported here are apparent. Closely spaced satellites, particularly around the centrosome, can also merge into single objects in 2D projections, limiting measurements of number, size and pericentrosomal clustering. These constraints are reduced when satellites form larger assemblies. For example, GFP-PCM1-N granules supported additional measurements such as circularity and sphericity, while SIM imaging of enlarged granules extended the measurable range. Unlike centrioles and cilia, satellites also have no defined dimensions or copy number that can serve as a ground truth. We therefore assessed SatelliteQ by comparing its measurements with independent methods and testing their robustness across segmentation parameters and biological conditions.

A further limitation concerns time-lapse analysis; because each frame is analyzed independently, SatelliteQ reports changes in satellite populations within cells rather than the trajectories of individual satellites. These limitations suggest clear directions for further development. Object tracking could resolve the behavior of individual satellites over time, while improved segmentation for densely packed cells and deep-learning approaches such as those recently applied to centrosome phenotyping by CenSegNet could extend satellite analysis to larger and more complex samples (Cheng et al., 2026). Incorporating reference structures beyond the centrosome and cilium, such as microtubules, would further expand the spatial questions that SatelliteQ can address.

SatelliteQ opens questions that could not previously be asked quantitatively; how satellite properties vary across tissues, differentiation states and physiological contexts, how they respond to the growing list of satellite-linked disease genes, and how they relate, cell by cell, to the centrosome and cilium phenotypes measured by existing tools. More broadly, the object-based, multiparametric approach established here offers a strategy for quantifying other membrane-less organelles whose properties no single measurement can capture.

## MATERIALS and METHODS

### Software and code availability

SatelliteQ is open-source software released under the MIT license, available at GitHub (https://github.com/Cytolab-Koc/SatelliteQ). The repository contains the Fiji macro, the Python module, an example input image, the settings template and a user guide. The Fiji macro was written for Fiji (ImageJ 1.54u8) and uses standard Fiji functions (Bio-Formats import, channel duplication, Z Project, Subtract Background, thresholding, Analyze Particles, Auto Local Threshold, binary Watershed, ROI Manager, Skeletonize, AnalyzeSkeleton and Measure). Three-dimensional analysis requires the 3D Objects Counter+ plugin (v0.1.0), which is installed through its Fiji update site. The Python module was written for Python 3.14.7 and uses NumPy, pandas, Matplotlib, Tkinter, openpyxl and Plotly, all of which are free and installed with pip.

### SatelliteQ image analysis pipeline

SatelliteQ accepts fluorescence microscopy images that contain at least one satellite channel; additional channels can contain a centrosome marker, a membrane or cytoplasmic marker for cell segmentation, or a primary cilium marker. Images are analyzed in one of four modes, 2D single image, 3D z-stack or time-lapses in either format. In 2D mode the macro generates a maximum intensity projection of the z-stack and all measurements are made on the projection; in 3D mode the z-stack is retained and objects are segmented across all sections. When calibration information is present in the image metadata, the physical pixel size and, for 3D images, the distance between consecutive z-sections are read and used to convert all measurements into micrometers; x and y positions are the pixel coordinates multiplied by the pixel size, and z positions are the section number multiplied by the z-step. Calibration values are carried through the analysis and written to the output.

The satellite channel is duplicated so that the raw data are not modified, and background fluorescence is reduced with Fiji’s rolling-ball Subtract Background function using a user-defined radius chosen according to satellite size and the amount of slowly varying background in the image. In 3D mode the same subtraction is applied to every section of the stack. The background-corrected image is then thresholded to generate a binary satellite mask. SatelliteQ offers the automatic thresholding methods Otsu, Triangle, Moments, Default, Li and Yen, a fixed manual threshold and a hysteresis-like option; in 3D mode, automatic thresholding uses only Otsu, because the other automatic methods were unstable on z-stacks, with a fixed manual threshold available as an alternative. Because satellite intensity and background differ between cell types, markers, treatments and microscopes, the threshold method and rolling-ball radius are chosen once on a test image and then applied to all images in the experiment; both values are recorded in the output (Table S2).

In 2D mode, objects in the binary mask are identified with Analyze Particles. User-defined minimum and maximum size limits remove small foreground regions that represent noise and, when required, unusually large regions; an optional binary Watershed step separates touching objects. An optional large-cluster splitting mode can further separate large 2D objects above a user-defined size using local intensity peaks. In 3D mode, connected voxels in the thresholded stack are identified with 3D Objects Counter+, which receives the selected threshold and size limits from the macro; objects touching the stack borders can optionally be excluded. In both modes each accepted object is stored as an ROI, assigned a unique Satellite ID and measured for position, boundary, size and fluorescence intensity.

When a cell boundary channel is provided, cell regions can be segmented from membrane, diffuse cytoplasmic or cytoskeletal signal or can be defined manually in the ROI editor. When no boundary marker is available, cell regions can instead be approximated as centrosome-centered circles in 2D or spheres in 3D with a user-defined radius; per-cell cilium status is reported only for marker-based or manually defined cell regions. Automated cell segmentation can use a global automatic threshold, a local automatic threshold or a fixed manual threshold; global methods include Otsu, Triangle, Moments, Default, Li, Yen and Huang, and local methods include Phansalkar, Sauvola, Bernsen, Niblack, Mean and Median with a user-defined local radius.

When a centrosome channel is provided, centrosomes can be detected automatically as bright puncta on a maximum-intensity projection after background subtraction and smoothing, using a percentage-of-maximum intensity threshold or a manual intensity threshold, or can be selected manually in the ROI editor. Detected centrosomes are displayed for review and their coordinates are stored as reference positions. Satellites and centrosomes are then assigned to cells.

Cells with more than one detected centrosome, for example cells with duplicated, separated or amplified centrosomes, are handled by one of three user-selected rules. Each satellite is assigned to the nearest centrosome in the same cell, the midpoint between the centrosomes is used as the reference position, or the cell is excluded from centrosome-dependent measurements. Cells with a single centrosome are analyzed normally.

After segmentation, all detected satellites are displayed as numbered objects in the ROI editor for visual inspection. In 2D analyses, including time-lapse data, incorrect detections can be deleted, ROI boundaries edited and missed satellites added before quantification, and segmentation can be repeated with different threshold, background-subtraction or size settings. In 3D analyses, review is visual; detected objects are displayed in a scrollable z-stack and the user either accepts the detections or repeats segmentation with different settings, without editing individual 3D objects. Images of the detected objects can be saved for quality control, and all user-defined settings are written to the output file so that the analysis can be reproduced.

For time-lapse analysis, the user selects the first and last frames and the time interval between frames, and the 2D or 3D workflow is applied to each frame with the same segmentation settings. Frame number and elapsed time are recorded for every measurement. Frames are analyzed independently, so SatelliteQ measures changes in satellite number, size, intensity, position and spatial distribution over time but does not track individual satellites between frames.

### Cilium detection

When a primary cilium channel is provided and cilium analysis is activated, the cilium channel is analyzed as a maximum-intensity projection. Background is reduced with a user-defined rolling-ball radius and the signal is smoothed with a Gaussian filter and thresholded; by default the threshold is set to the image mean plus two standard deviations, with percentage-of-maximum and fixed manual thresholds available as alternatives. Thresholded objects are identified with Analyze Particles and filtered according to their skeleton path length. For each accepted cilium, the thresholded signal is skeletonized and its length is measured with AnalyzeSkeleton (v3.4.2). The default cilium-length range is 1 to 10 µm and can be changed by the user; the minimum accepted length also defines the ciliation threshold. Cilium candidates are displayed for review, and incorrect detections can be deleted. Cilium count and length values are written to separate Fiji output files and incorporated into the Python report. Cells without an accepted cilium are classified as non-ciliated. Cilium analysis is supported for 2D analyses and for 3D datasets through maximum-intensity projection. For the comparison of ciliated and unciliated cells (Fig. S7D–I), serum-starved RPE1 cells were classified by ARL13B staining with the cilium detection module.

### Quantification of satellite properties

Satellite size is reported as area in 2D mode, calculated from the number of pixels inside each object and the calibrated pixel size, and as apparent volume in 3D mode, calculated from the number of voxels together with the pixel size and z-step. Fluorescence intensity is measured on the background-corrected grayscale image within the boundary defined by the mask. For each satellite, integrated intensity is the sum of all pixel or voxel intensities inside the object and mean intensity is their average; integrated intensities are reported as raw, uncalibrated sums. Because these values depend on acquisition and sampling settings, intensities are compared only within experiments acquired with identical settings, and control-normalized values are used when comparisons span experiments. At the cell level, total satellite intensity is the sum of the integrated intensities of all satellites assigned to that cell and mean satellite intensity is their average; total and mean satellite size are calculated in the same way.

The straight-line distance between the centroid of each satellite and its assigned centrosome is calculated from x and y positions in 2D mode and from x, y and z positions in 3D mode. When nearest-centrosome assignment is selected, the closest centrosome inside the same cell is used. The mean satellite-to-centrosome distance is calculated for each cell from all satellites with a valid centrosome assignment.

Spatial entropy describes how evenly satellite fluorescence is distributed within a cell and is calculated for each cell and, in time-lapse data, for each frame. The cell domain is the segmented cell boundary when available and otherwise a circle of equal calibrated cell area. The domain is divided into 5 × 5, 10 × 10 and 20 × 20 grids, and for each grid the Shannon entropy H = −Σ pⱼ ln pⱼ is calculated over the bins inside the domain, where pⱼ is the fraction of the total integrated satellite intensity in bin j. Each value is normalized by ln(min(K, N)), where K is the number of bins inside the domain and N the number of satellites in the cell, so that normalized entropy ranges from 0, all signal in one bin, to 1, signal spread uniformly across the cell. The reported spatial entropy is the mean of the three normalized values; the individual grid values and a count-based variant are retained in the output for quality control.

To quantify the enrichment of satellites around the centrosome, SatelliteQ calculates a pericentrosomal clustering index (PCI), following intensity-based clustering measurements used previously (e.g. Aydin et al., 2020). The integrated satellite intensity within a circle (2D) or sphere (3D) of radius r₁ centered on the centrosome is divided by the integrated satellite intensity in the surrounding annulus or shell between r₁ and r₂, with r₁ = 3 µm and r₂ = 12 µm. Radial distance is calculated in x-y for 2D analyses and in x-y-z for 3D analyses. The inner 0 to 3 µm intensity, outer 3 to 12 µm intensity and total 0 to 12 µm intensity are also reported separately. PCI increases when satellites cluster around the centrosome, decreases when they disperse, and is reported as undefined when the outer region contains no satellite signal.

Circularity is reported for 2D objects as measured by Analyze Particles, 4π × area / perimeter². Sphericity of 3D objects is taken from the 3D Objects Counter+ output when reported and is otherwise calculated as π(1/3)(6V)(2/3)/S from the volume V and surface area S of the object; the source of each value is recorded in the output. Both values approach 1 for round objects.

### Data processing, reports and interactive viewers

The Python module analyzes one Fiji output file or a folder containing several files, reads the tables and converts the column names into standardized SatelliteQ names. Area and volume, and cell area and cell volume, are kept as separate measurements. Each satellite retains its image, Satellite ID, Cell ID, experimental condition, replicate and, where applicable, frame. Condition and replicate are read from the folder names defined by the user. When several files are combined, unique identifiers are generated for each image, cell and satellite so that objects from different images that share the same numerical ID are never merged.

The Python module writes a single Excel report with separate sheets for satellite-level, cell-level, image-level and, for time-lapse data, frame-level measurements; every row is labeled with its condition, replicate, image, frame, cell and satellite identifiers, and the analysis settings are written to a separate sheet. Two interactive viewers are generated with Plotly and opened in the web browser. The Graph Explorer plots any measurement across conditions or replicates as point, cell-panel, bar, box, line and histogram plots, and can display means, standard deviations, sample numbers, outlier annotations and x-y comparisons with linear fits and R² values. The Experiment Explorer displays cell-level measurements by condition and replicate, with optional mean ± SD, normalization and statistical-comparison overlays. Values can be normalized to a reference condition; the mean of each replicate of the reference condition is set to 1 and all values from the corresponding replicate are divided by that mean. The Spatial Viewer reconstructs the measured spatial organization of each source image; in 2D it displays satellite and centrosome x-y coordinates together with available cell boundaries, and in 3D, when valid z coordinates are available, satellites and centrosomes are displayed at their measured x-y-z positions with sphere-based estimated cell regions.

### Image analysis settings and performance

All images were analyzed with SatelliteQ using the mode, threshold method, rolling-ball radius and size limits listed for each experiment in Table S2. SatelliteQ runs on standard personal computers that support Fiji and Python, for example systems with an Intel Core i5/i7 or Apple M1 processor and 8 to 16 GB RAM. For representative 1,024 × 1,024-pixel confocal images, automated 2D analysis typically takes approximately 5 to 20 s per image, and 3D analysis of z-stacks of 20 to 30 optical sections approximately 40 to 180 s per stack, with peak memory use of approximately 0.5 to 1.5 GB in 2D and 1.5 to 4 GB in 3D. These times cover image processing, segmentation, quantification and export of results and exclude manual review. Actual performance depends on image dimensions, z-stack depth, satellite density and segmentation settings.

### Analysis in Imaris

Three-dimensional quantification of centriolar satellites was performed with Imaris version 10.2.0 (Oxford Instruments). Confocal z-stacks were imported without projection and PCM1-positive satellites were segmented in three dimensions with the Imaris Surfaces module. A threshold value of 5000, the same value used for SatelliteQ, was selected on representative images and kept constant for all images. Satellite number was defined as the number of segmented surfaces within each cell. Individual satellite volume was obtained directly from the Imaris surface measurements and total satellite volume was calculated as the sum of the volumes of all satellites assigned to a cell; mean satellite volume is total volume divided by satellite number. Integrated fluorescence intensity was obtained from the intensity-sum measurement of each segmented PCM1 surface and total satellite fluorescence intensity per cell was calculated as the sum of the intensity values of all satellites within that cell. Centrosomes were identified with centrin2 and satellite-to-centrosome distances were calculated from individual satellites to the centrosome reference point.

### Cell culture

Human hTERT-RPE1, HeLa, HEK293T and U2OS cells were used for validation of SatelliteQ and for comparison of satellite properties across cell types. RPE1 cells were cultured in DMEM/F12 and HeLa, U2OS and HEK293T cells in DMEM, each supplemented with 10% fetal bovine serum (FBS) and penicillin-streptomycin. Cells were maintained at 37 °C in a humidified incubator with 5% CO₂ and tested regularly for mycoplasma contamination. For comparison of satellite properties between cell types, cells were grown on glass coverslips, fixed at 80 to 90% confluency, stained for PCM1, α-tubulin and centrin and imaged with identical settings within each experiment.

RPE1 PCM1 knockout cells carrying doxycycline-inducible GFP-PCM1-N were used to follow de novo granule formation. (Begar et al., 2026). Expression was induced with 1 µM doxycycline; for live imaging, doxycycline was added at the start of acquisition and the first frame with detectable GFP-PCM1-N granules was defined as the beginning of the granule formation time course. Control and CEP290- or NPHP1-mutant urine-derived renal epithelial cells (URECs) were established and conditionally immortalized as described (Garcia et al., 2022; Malglaive et al., 2025; Menguy et al., 2026). In brief, cells isolated from urine were transduced with a thermosensitive SV40 large T antigen and expanded at the permissive temperature of 33 °C and 5% CO₂ in UREC proliferation medium consisting of REBM basal medium (CC-3191, Lonza) with 1× REGM SingleQuots supplements (CC-4127, Lonza), 10 ng/ml recombinant human EGF (R&D Systems) and 2% certified FBS (Invitrogen). For ciliogenesis assays, 400,000 URECs were seeded onto coverslips in 24-well plates and shifted to the non-permissive temperature of 39 °C for 3 days to arrest proliferation and induce ciliogenesis before fixation.

### Treatments and siRNA transfection

RPE1 cells were treated with 5 µg/ml nocodazole or an equal volume of DMSO for 1 h before fixation. For disruption of weak hydrophobic interactions, RPE1 cells were treated with 5% 1,6-hexanediol for 5 min before fixation; DMSO-treated cells processed in parallel served as controls. For depletion of CEP290, a previously characterized siRNA (GAUACUCGGUUUUUACGUAUUUU; Dharmacon) and control siRNA#1 (D-001210-02-05, Dharmacon) were used. Cells seeded on coverslips at 70% confluency were transfected twice sequentially with 25 nM siRNA using Lipofectamine RNAiMAX (13778- 150, Life Technologies) in Opti-MEM (31985062, Life Technologies) according to the manufacturer’s instructions, and depletion was confirmed by immunofluorescence 48 h after transfection. For ciliation analysis, transfected cells were serum starved for 24 h before fixation and ciliation was scored by ARL13B staining. For the comparison of ciliated and unciliated cells (Fig. S7C–I), untransfected RPE1 cells were serum starved for 24 h before fixation.

### Immunofluorescence and antibodies

Cells were fixed in ice-cold methanol at −20 °C for 10 min, rehydrated in PBS and blocked in 3% BSA (Sigma-Aldrich) in PBS with 0.1% Triton X-100 for 30 min. Coverslips were incubated with primary antibodies in blocking solution for 1 h at room temperature, followed by Alexa Fluor 488-, 594- or 633-conjugated secondary antibodies for 1 h at room temperature, and mounted in Mowiol containing N-propyl gallate (Sigma-Aldrich). Antibodies used in this study are described in Table S3.

### Microscopy

Fixed cells were imaged at room temperature on a Leica Stellaris 8 inverted laser-scanning confocal microscope equipped with 405, 488, 561 and 633 nm laser lines, one photomultiplier tube and three hybrid detectors, using HC PL APO CS2 63×/1.4 NA or 40×/1.3 NA oil-immersion objectives with the pinhole set to 1 Airy unit. Z-stacks of 15 to 30 optical sections at 0.3 to 0.5 µm spacing were acquired at 1,024 × 1,024 pixels in unidirectional xyz scan mode at 400 to 600 Hz with LAS X software. Live cells were imaged on the same microscope with the HC PL APO CS2 63×/1.4 NA objective and an incubation chamber maintained at 37 °C and 5% CO₂; cells were imaged every 1.5 min for GFP-PCM1-N assembly experiments. Structured illumination microscopy (SIM) was performed on an Elyra 7 microscope equipped with Lattice SIM (Carl Zeiss Microscopy), a Plan-Apochromat 63×/1.4 NA oil objective and 405, 488, 561 and 633 nm laser lines. Images were acquired with an sCMOS camera (version 4.2 CL HS) with a z-step of 110 nm across a total imaging depth of 5 to 10 µm, and SIM reconstruction was performed with ZEN Black software. Reconstructed images of GFP-PCM1-N assemblies were analyzed with SatelliteQ to quantify granule circularity. Maximum intensity projection, 3D rendering and brightness and contrast adjustments applied after acquisition were identical for all samples within an experiment.

### Statistical analysis

Statistical analyses and graph generation were performed with GraphPad Prism version 8. The number of independent biological replicates and sample sizes are indicated in the corresponding figure legends. Data are presented as mean ± SD. The statistical test used for each comparison is indicated in the corresponding figure legend; paired two-tailed Student’s t tests were used when the same cells were measured in both analysis modes, unpaired two-tailed Student’s t tests for two-group comparisons of different cells, and one-way ANOVA with multiple comparisons. Statistical significance was defined as P < 0.05; *, P < 0.05; **, P < 0.01; ***, P < 0.001; ****, P < 0.0001; ns, not significant.

### Ethics

UREC lines were derived from urine samples collected from individuals who provided written informed consent and were anonymized in accordance with the NPH1 protocol, approved by the French National Committee for the Protection of Persons (ID-RCB no. 2016-A00541-50), and the RIM-C’IL-LICO protocol, approved by the CPP Île-de-France IV (ID-RCB 2020-A02356-33). The research was conducted in accordance with the Declaration of Helsinki on human experimentation of the World Medical Association. The UREC lines are included in the Institut Imagine Biocollection declared to the French Ministry of Research (DC-2024-6350).

## ACKNOWLEDGEMENTS

We thank the members of the Firat-Karalar laboratory for helpful discussions and critical reading of the manuscript. We thank Dila Gülensoy, Serra Doğanata, Halil Bayraktar, and Çiğdem Gündüz Demir for their valuable contributions to the conceptualization of questions related to quantification of centriolar satellite properties. We are grateful to Dr. Marc Fila (CHU de Montpellier) and Jean-Michel Rozet (Laboratory of Genetics in Ophthalmology, Imagine Institute) for their contributions to the patient-derived UREC studies and for providing patient-derived UREC cells. We also sincerely thank the patients and their families for their participation and contribution to this research. We thank Ronald Roepman (Radboud University Medical Center, Nijmegen) and his group for discussions on ciliopathy phenotyping and also other members of the ERDERA PREDICT consortium. This work was supported by the European Research Council Starting Grant Satellite Homeostasis under Grant Agreement 101078097, TUBITAK Grant 123N048 through the ERDERA joint transnational call project PREDICT, and Istanbul Development Agency YEP-0057 Grant, all awarded to E.N. Firat-Karalar. Work in the Saunier laboratory was supported by the European Union’s Horizon 2020 programme (EJP RD, PREDICT, Grant Agreement No. 825575) and Horizon Europe programme (TheRaCil, Grant Agreement No. 101080717).

During the preparation of this work, Efe Begar used OpenAI ChatGPT to assist with the development, debugging, and refinement of the Fiji macro and Python code used in the SatelliteQ workflow. The authors reviewed, tested and carefully edited the AI-assisted code as needed and take full responsibility for the functionality of the software and the content of the published article.

