## Supplementary Figures for "SatelliteQ: an open-source workflow for multidimensional, object-based quantification of centriolar satellites"

### Figure S1

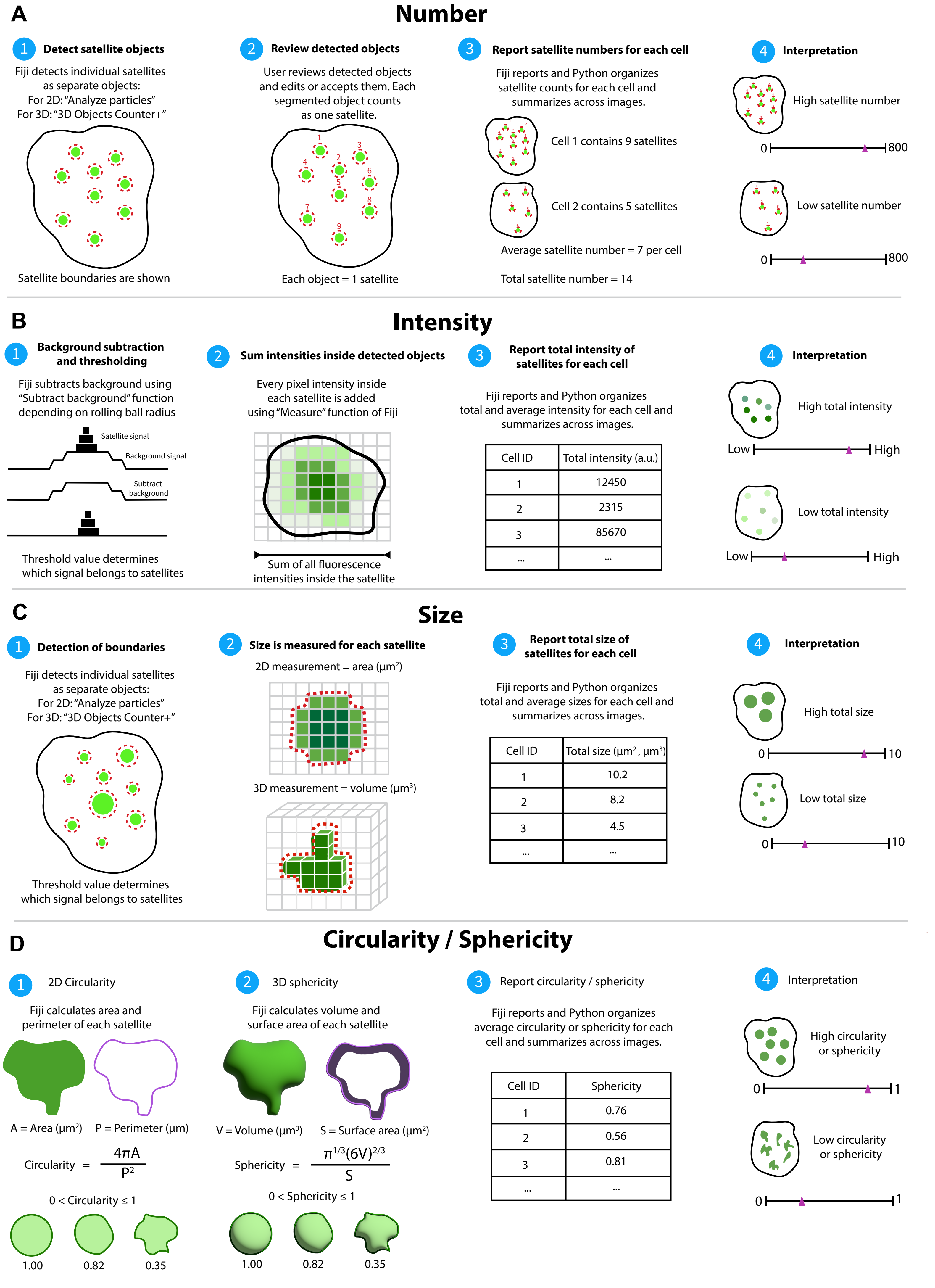
A
Number
1 Detect satellite objects
Fiji detects individual satellites as separate objects:
For 2D: "Analyze particles"
For 3D: "3D Objects Counter+"
Satellite boundaries are shown
2 Review detected objects
User reviews detected objects and edits or accepts them. Each segmented object counts as one satellite.
Each object = 1 satellite
3 Report satellite numbers for each cell
Fiji reports and Python organizes satellite counts for each cell and summarizes across images.
Cell 1 contains 9 satellites
Cell 2 contains 5 satellites
Average satellite number = 7 per cell
Total satellite number = 14
4 Interpretation
High satellite number
0
800
Low satellite number
0
800
B
Intensity
1 Background subtraction and thresholding
Fiji subtracts background using "Subtract background" function depending on rolling ball radius
Satellite signal
Background signal
Subtract background
Threshold value determines which signal belongs to satellites
2 Sum intensities inside detected objects
Every pixel intensity inside each satellite is added using "Measure" function of Fiji
Sum of all fluorescence intensities inside the satellite
3 Report total intensity of satellites for each cell
Fiji reports and Python organizes total and average intensity for each cell and summarizes across images.
Cell ID | Total intensity (a.u.)
1 | 12450
2 | 2315
3 | 85670
... | ...
4 Interpretation
High total intensity
Low
High
Low total intensity
Low
High
C
Size
1 Detection of boundaries
Fiji detects individual satellites as separate objects:
For 2D: "Analyze particles"
For 3D: "3D Objects Counter+"
Threshold value determines which signal belongs to satellites
2 Size is measured for each satellite
2D measurement = area (μm²)
3D measurement = volume (μm³)
3 Report total size of satellites for each cell
Fiji reports and Python organizes total and average sizes for each cell and summarizes across images.
Cell ID | Total size (μm², μm³)
1 | 10.2
2 | 8.2
3 | 4.5
... | ...
4 Interpretation
High total size
0
10
Low total size
0
10
D
Circularity / Sphericity
1 2D Circularity
Fiji calculates area and perimeter of each satellite
A = Area (μm²)
P = Perimeter (μm)
Circularity = 4πA / P²
0 < Circularity ≤ 1
1.00
0.82
0.35
2 3D sphericity
Fiji calculates volume and surface area of each satellite
V = Volume (μm³)
S = Surface area (μm²)
Sphericity = π^(1/3)(6V)^(2/3) / S
0 < Sphericity ≤ 1
1.00
0.82
0.35
3 Report circularity / sphericity
Fiji reports and Python organizes average circularity or sphericity for each cell and summarizes across images.
Cell ID | Sphericity
1 | 0.76
2 | 0.56
3 | 0.81
... | ...
4 Interpretation
High circularity or sphericity
0
1
Low circularity or sphericity
0
1

**Supplementary Figure 1. SatelliteQ quantifies satellite number, fluorescence intensity, size and shape.**

**(A)** Schematic representation of satellite number quantification. Individual satellites are segmented with Analyze Particles in 2D or 3D Objects Counter+ in 3D, displayed in the ROI editor for review and editing, and each accepted object receives a unique Satellite ID and contributes one object to the count. Counts are summarized by the Python module at the cell, image and experiment levels. Examples illustrate cells with relatively high or low satellite number. **(B)** Schematic representation of satellite fluorescence intensity quantification. Background fluorescence is reduced with Fiji's Subtract Background function using a user-defined rolling-ball radius, and thresholding defines the boundaries of satellite objects. Intensities are measured on the background-corrected grayscale image within each boundary. Integrated intensity is the sum of pixel or voxel intensities within each satellite, and cell-level totals and means are calculated by the Python module. Examples illustrate satellite populations with high or low total fluorescence intensity. **(C)** Schematic representation of satellite size quantification. Object boundaries defined during segmentation give satellite area in $\mu m^2$ in 2D analysis and apparent volume in $\mu m^3$ in 3D analysis, which are summarized as total and mean satellite size for each cell. Examples illustrate cells with relatively high or low total satellite size. **(D)** Schematic representation of shape measurements. Circularity is reported for 2D objects and sphericity for 3D objects, calculated from area and perimeter or volume and surface area; both approach 1 for round objects. Examples illustrate round satellites and the larger irregular granules for which these measurements are informative.

**Figure S2**

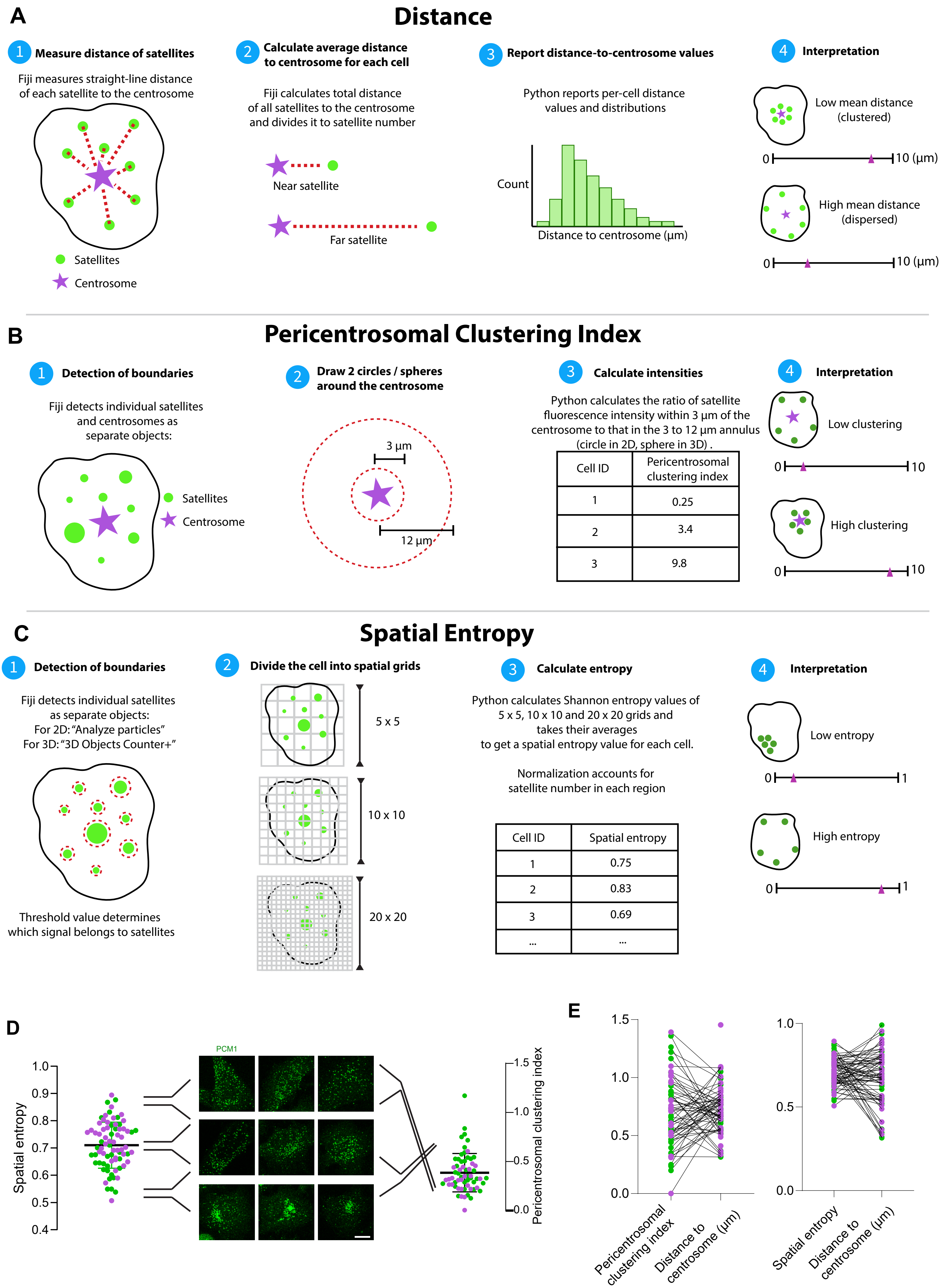

A
Distance
1 Measure distance of satellites
Fiji measures straight-line distance of each satellite to the centrosome
Satellites
Centrosome
2 Calculate average distance to centrosome for each cell
Fiji calculates total distance of all satellites to the centrosome and divides it to satellite number
Near satellite
Far satellite
3 Report distance-to-centrosome values
Python reports per-cell distance values and distributions
Count
Distance to centrosome (μm)
4 Interpretation
Low mean distance (clustered)
0
10 (μm)
High mean distance (dispersed)
0
10 (μm)
B
Pericentrosomal Clustering Index
1 Detection of boundaries
Fiji detects individual satellites and centrosomes as separate objects:
Satellites
Centrosome
2 Draw 2 circles / spheres around the centrosome
3 μm
12 μm
3 Calculate intensities
Python calculates the ratio of satellite fluorescence intensity within 3 μm of the centrosome to that in the 3 to 12 μm annulus (circle in 2D, sphere in 3D) .
Cell ID | Pericentrosomal clustering index
1 | 0.25
2 | 3.4
3 | 9.8
4 Interpretation
Low clustering
0
10
High clustering
0
10
C
Spatial Entropy
1 Detection of boundaries
Fiji detects individual satellites as separate objects:
For 2D: "Analyze particles"
For 3D: "3D Objects Counter+"
Threshold value determines which signal belongs to satellites
2 Divide the cell into spatial grids
5 x 5
10 x 10
20 x 20
3 Calculate entropy
Python calculates Shannon entropy values of 5 x 5, 10 x 10 and 20 x 20 grids and takes their averages to get a spatial entropy value for each cell.
Normalization accounts for satellite number in each region
Cell ID | Spatial entropy
1 | 0.75
2 | 0.83
3 | 0.69
... | ...
4 Interpretation
Low entropy
0
1
High entropy
0
1
D
Spatial entropy
1.0
0.9
0.8
0.7
0.6
0.5
0.4
PCM1
1.5
1.0
0.5
0.0
Pericentrosomal clustering index
E
1.5
1.0
0.5
0.0
Pericentrosomal clustering index
Distance to centrosome (μm)
1.0
0.5
0.0
Spatial entropy
Distance to centrosome (μm)

**Supplementary Figure 2. SatelliteQ quantifies satellite spatial organization through centrosome distance, PCI and spatial entropy.**

**(A)** Schematic representation of satellite-to-centrosome distance measurements. The straight-line distance between the centroid of each satellite and its assigned centrosome is calculated in 2D or 3D according to the analysis mode, and the mean distance is calculated for each cell. Examples illustrate satellite populations with low or high mean centrosome distance. **(B)** Schematic representation of the pericentrosomal clustering index (PCI). The integrated satellite intensity within a circle of radius 3 μm centered on the centrosome (sphere in 3D) is divided by the integrated satellite intensity in the surrounding region up to 12 μm. PCI increases when satellites cluster around the centrosome and decreases when they disperse. Examples illustrate satellite populations with high and low PCI. **(C)** Schematic representation of spatial entropy analysis. The cell region is divided into 5 × 5, 10 × 10 and 20 × 20 grids, the distribution of satellite fluorescence across grid cells is used to calculate normalized Shannon entropy, and the reported value is the mean of the three grids. Values approach 1 for uniformly distributed and 0 for concentrated satellite signal. **(D)** Spatial entropy and PCI per cell in RPE1 cells stained for PCM1. Images show representative cells at the indicated entropy ranges, from clustered to dispersed satellite distributions. Each dot represents one cell; colours indicate biological replicates (green and magenta). Black bars represent mean ± SD. Scale bar: 10 μm. **(E)** Per-cell comparison of the spatial organization measurements, PCI against mean satellite-to-centrosome distance and spatial entropy against mean distance; black lines connect the values of the same cell. Distance values are normalized so that the mean of each replicate equals the mean of the compared measurement. Results are from 2 biological replicates (more than 38 cells per replicate). Analysis settings are listed in Table S2.

### Figure S3

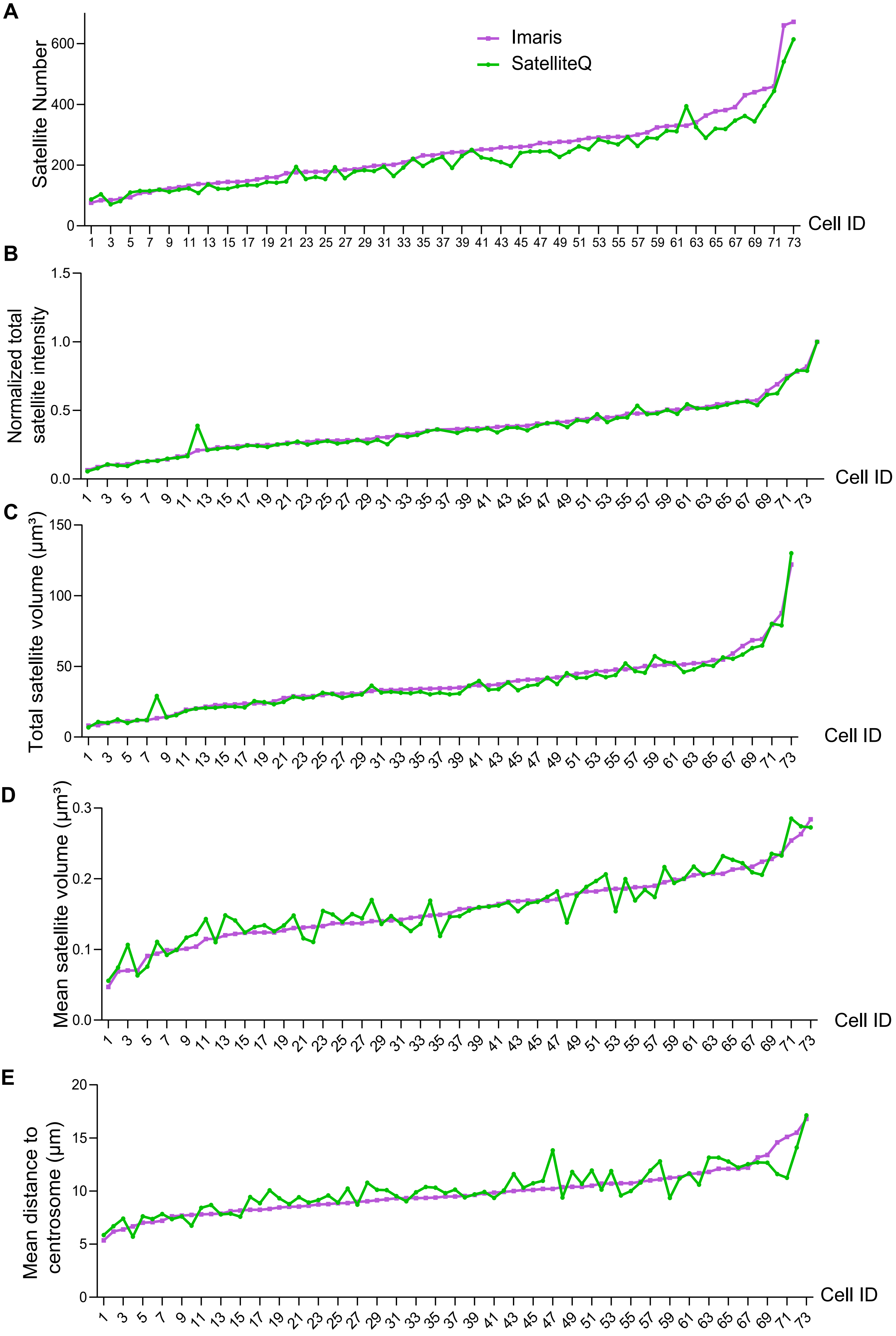
A
B
C
D
E
Imaris
SatelliteQ
Satellite Number
Normalized total satellite intensity
Total satellite volume (μm³)
Mean satellite volume (μm³)
Mean distance to centrosome (μm)
Cell ID

**Supplementary Figure 3. SatelliteQ reproduces cell-to-cell variation measured by Imaris.**

**(A to E)** The same 3D images of RPE1 cells stained for PCM1 were analyzed independently with SatelliteQ and Imaris, and the per-cell values from the two methods are overlaid, with cells ranked by their Imaris value in each panel, for satellite number **(A)**, total satellite fluorescence intensity **(B)**, total satellite volume **(C)**, mean satellite volume **(D)** and mean satellite-to-centrosome distance **(E)**. Purple, Imaris; green, SatelliteQ. Intensity values are normalized to the maximum value of each replicate, set to 1. The same images and thresholding settings were used for the SatelliteQ and Imaris analyses. Results are from 2 biological replicates (73 cells in total). Analysis settings are listed in Table S2.

### Figure S4

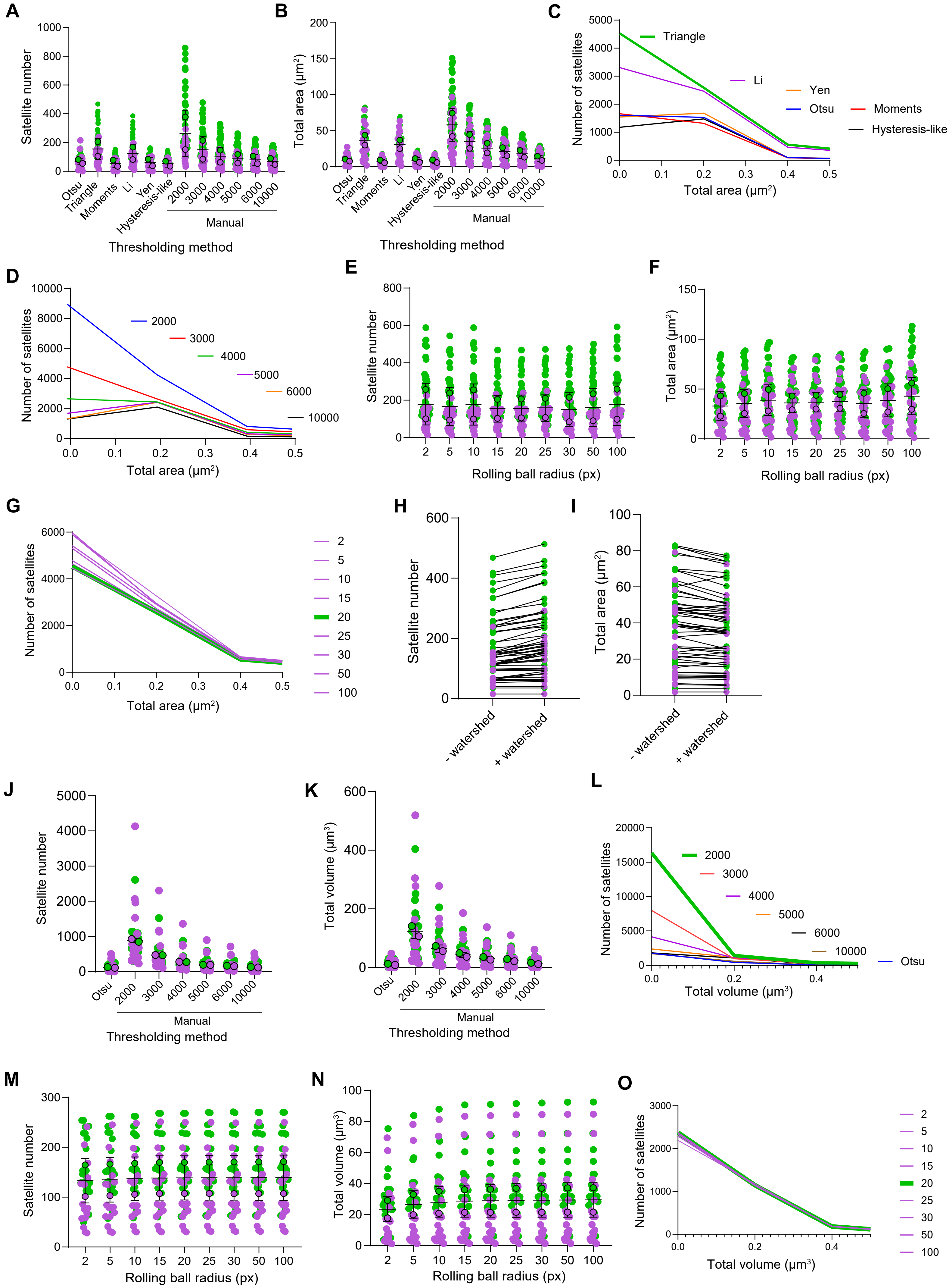
A
Satellite number
Otsu
Triangle
Moments
Li
Yen
Hysteresis-like
2000
3000
4000
5000
6000
10000
Manual
Thresholding method
B
Total area (μm²)
Thresholding method
C
Number of satellites
Total area (μm²)
Triangle
Li
Yen
Otsu
Moments
Hysteresis-like
D
Number of satellites
Total area (μm²)
E
Satellite number
Rolling ball radius (px)
F
Total area (μm²)
Rolling ball radius (px)
G
Number of satellites
Total area (μm²)
H
Satellite number
- watershed
+ watershed
I
Total area (μm²)
J
Satellite number
Thresholding method
K
Total volume (μm³)
Thresholding method
L
Number of satellites
Total volume (μm³)
M
Satellite number
Rolling ball radius (px)
N
Total volume (μm³)
Rolling ball radius (px)
O
Number of satellites
Total volume (μm³)

**Supplementary Figure 4. Segmentation parameters affect absolute satellite measurements in 2D and 3D.**

Six images of RPE1 cells stained for PCM1, each containing multiple cells, were analyzed with one segmentation parameter varied at a time; all other settings were held at the values used in this study (Table S2). **(A and B)** Satellite number **(A)** and total satellite area **(B)** per cell for the indicated thresholding methods in 2D. **(C and D)** Size distribution of objects detected with each automatic thresholding method **(C)** and each manual threshold value **(D)** in 2D (bin width, 0.2 $\mu m^2$). **(E and F)** Satellite number **(E)** and total satellite area **(F)** per cell for rolling-ball radii of 2–100 pixels in 2D. **(G)** Size distribution of detected objects for each rolling-ball radius in 2D. **(H and I)** Satellite number **(H)** and total satellite area **(I)** per cell without and with watershed separation; lines connect measurements from the same cell. Watershed separation was applied only to the 2D workflow. **(J and K)** Satellite number **(J)** and total satellite volume **(K)** per cell for Otsu and the indicated manual threshold values in 3D. **(L)** Size distribution of detected objects for each threshold in 3D (bin width, 0.2 $\mu m^3$). **(M and N)** Satellite number **(M)** and total satellite volume **(N)** per cell for rolling-ball radii of 2–100 pixels in 3D. **(O)** Size distribution of detected objects for each rolling-ball radius in 3D. Each dot represents one cell. Colours indicate biological replicates, and green indicates the settings used for the corresponding analyses in this study. Black bars represent mean ± SD. Results are from 2 biological replicates (>24 cells per replicate). No statistical tests were applied.

**Figure S5**

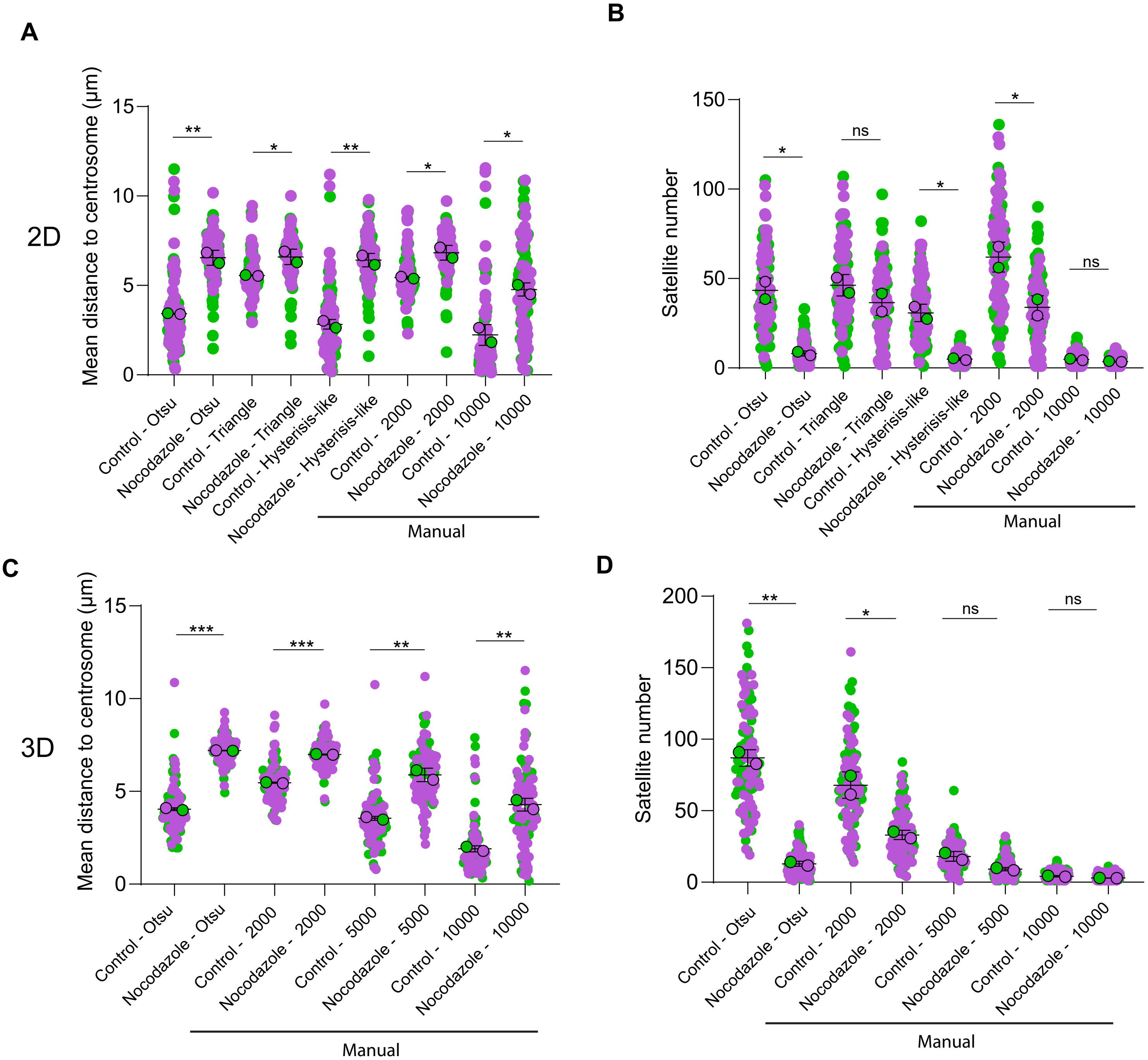
A
B
2D
Mean distance to centrosome (μm)
15
10
5
0
**
*
**
*
*
Control - Otsu
Nocodazole - Otsu
Control - Triangle
Nocodazole - Triangle
Control - Hysterisis-like
Nocodazole - Hysterisis-like
Control - 2000
Nocodazole - 2000
Control - 10000
Nocodazole - 10000
Manual
Satellite number
150
100
50
0
*
ns
*
*
ns
C
D
3D
***
***
**
**
Control - 5000
Nocodazole - 5000
200
150
100
50
0
**
*
ns
ns

**Supplementary Figure 5. Nocodazole-induced satellite phenotypes are detected across thresholding methods.**

Control and nocodazole-treated RPE1 cells stained for PCM1 were analyzed with the indicated thresholding methods, with each method applied identically to both conditions. **(A and B)** Mean satellite-to-centrosome distance **(A)** and satellite number **(B)** per cell in 2D. **(C and D)** The same measurements in 3D. Each dot represents one cell; colours indicate biological replicates (green and magenta) and larger outlined circles represent replicate means. Black bars represent mean ± SD. Results are from 2 biological replicates (more than 52 cells per replicate). Statistical significance was determined by two-tailed unpaired Student's t test ; *, $P < 0.05$; **, $P < 0.01$; ***, $P < 0.001$; ns, not significant. Analysis settings are listed in Table S2.

### Figure S6

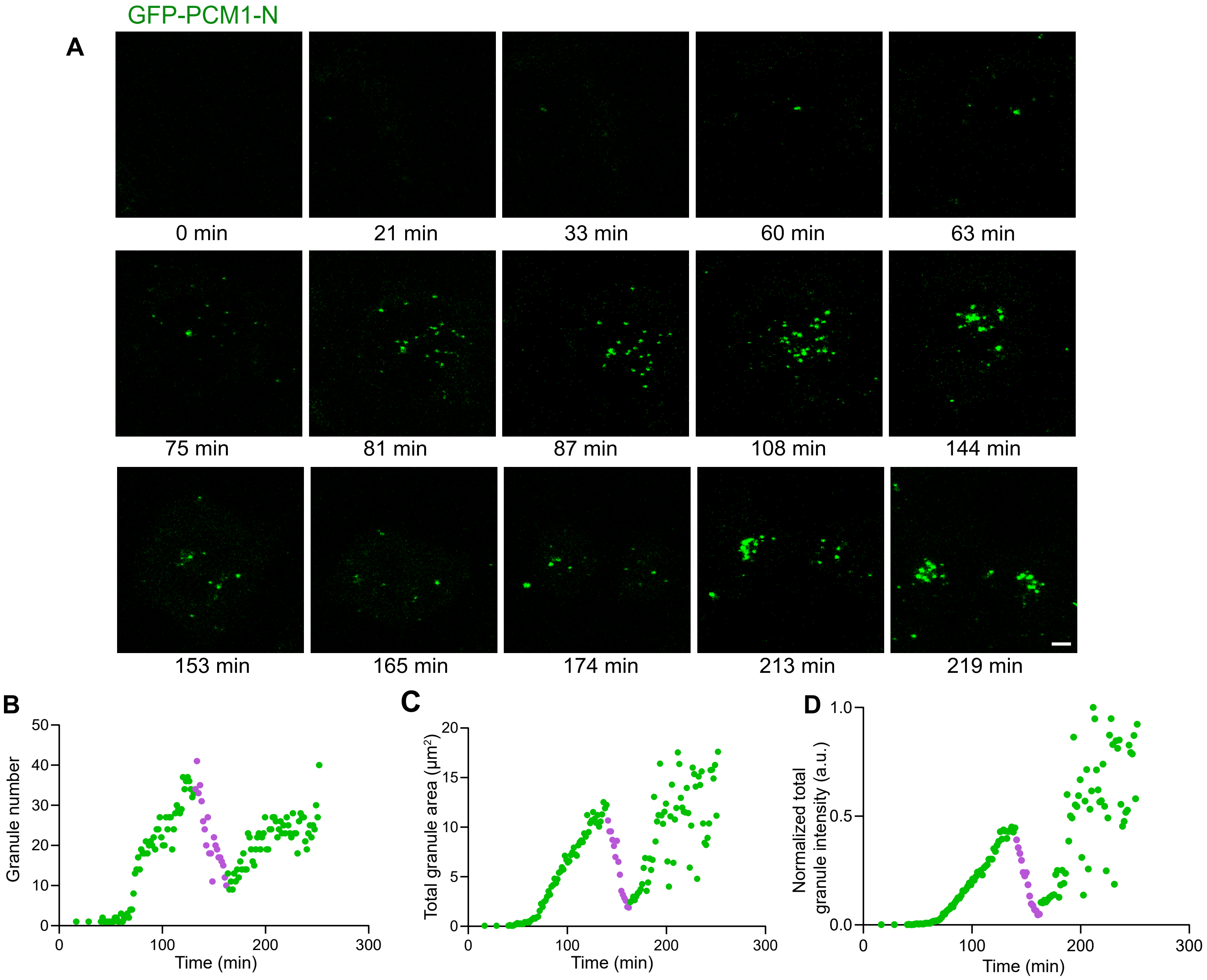

GFP-PCM1-N
A
0 min
21 min
33 min
60 min
63 min
75 min
81 min
87 min
108 min
144 min
153 min
165 min
174 min
213 min
219 min
B
Granule number
Time (min)
C
Total granule area (μm²)
Time (min)
D
Normalized total granule intensity (a.u.)
Time (min)

**Supplementary Figure 6. Single-cell dynamics of GFP-PCM1-N granule formation and remodeling.**

**(A)** Stills from time-lapse imaging of an RPE1 PCM1 knockout cell expressing doxycycline-inducible GFP-PCM1-N progressing through mitosis; times are relative to doxycycline addition and are indicated below each still. **(B to D)** Granule number **(B)**, total granule area **(C)** and normalized total granule fluorescence intensity **(D)** over time for the same cell. Magenta points represent measurements during mitosis.

### Figure S7

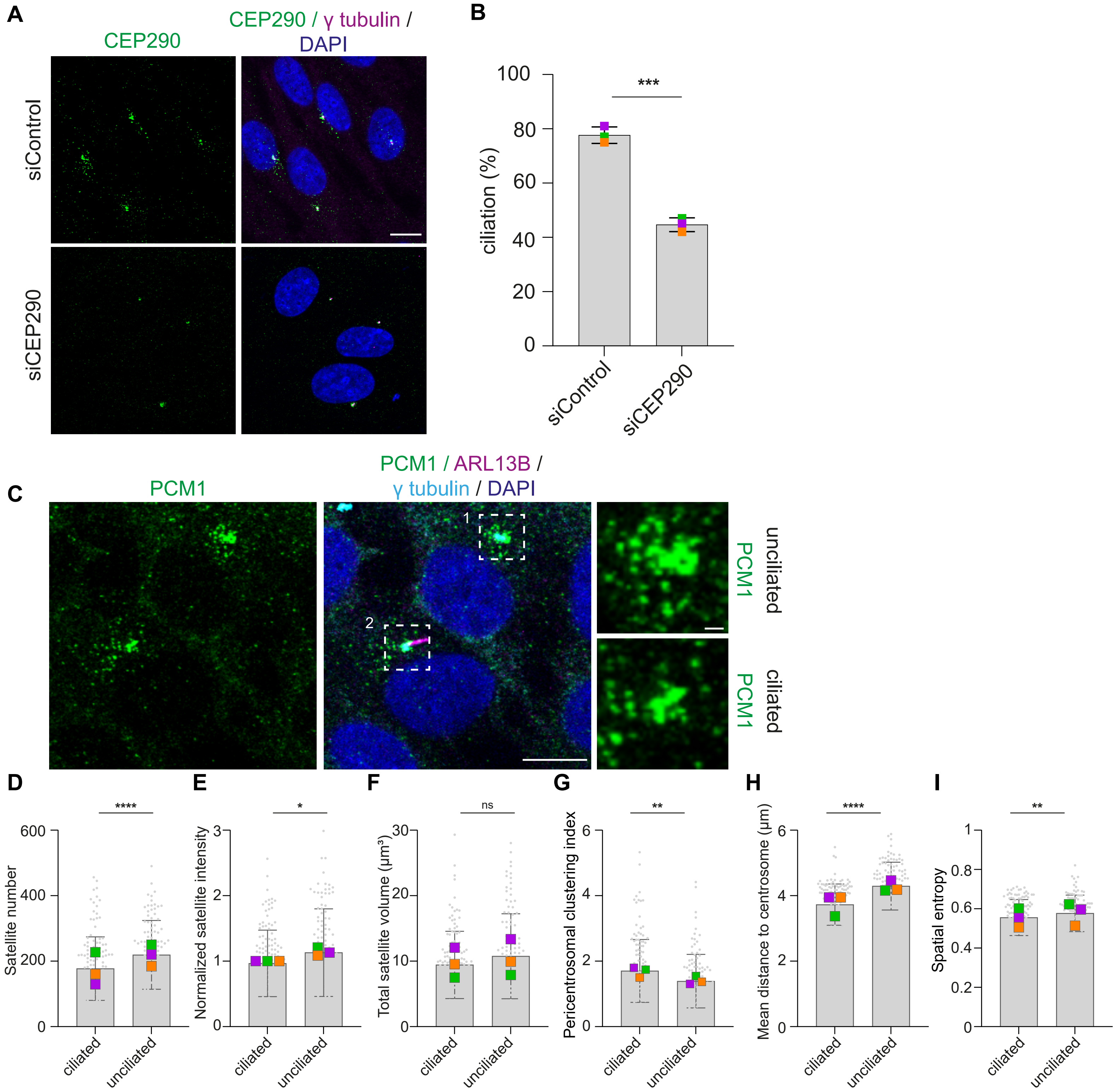
A
CEP290
CEP290 / γ tubulin / DAPI
siControl
siCEP290
B
ciliation (%)
***
siControl
siCEP290
C
PCM1
PCM1 / ARL13B / γ tubulin / DAPI
1
2
PCM1 unciliated
PCM1 ciliated
D
Satellite number
****
E
Normalized satellite intensity
*
F
Total satellite volume (µm³)
ns
G
Pericentrosomal clustering index
**
H
Mean distance to centrosome (µm)
****
I
Spatial entropy
**
ciliated
unciliated

**Supplementary Figure 7. Validation of CEP290 depletion and reduced ciliation in RPE1 cells.**

**(A)** Representative images of RPE1 cells transfected with control or CEP290 siRNA, serum starved for 24 h, and stained for CEP290 (green), γ-tubulin (magenta) and DAPI (blue). **(B)** Percentage of ciliated cells, scored by ARL13B staining. Coloured squares represent biological replicates; bars represent mean ± SD. Results are from 3 biological replicates (minimum 100 cells per replicate). **(C)** Representative images of RPE1 cells transfected with control siRNA, serum starved for 24 h and stained for PCM1 (green), ARL13B (magenta), γ-tubulin (cyan) and DAPI (blue). Boxed regions are shown at higher magnification and illustrate PCM1 organization in unciliated (1) and ciliated (2) cells. **(D to I)** Satellite number **(D)**, normalized total satellite fluorescence intensity **(E)**, total satellite volume **(F)**, pericentrosomal clustering index **(G)**, mean satellite-to-centrosome distance **(H)** and spatial entropy **(I)** in ciliated and unciliated RPE1 cells, quantified using the 3D workflow. Intensity values are normalized to the mean of the ciliated condition, set to 1. Each dot represents one cell; coloured squares represent replicate means. Bars represent mean ± SD. Results are from 3 biological replicates (minimum 100 cells per condition). Statistical significance was determined by unpaired two-tailed Student's $t$ test; *, $P < 0.05$; **, $P < 0.01$; ***, $P < 0.001$; ns, not significant.. Scale bar: 10 µm. Scale bar for inset: 1 µm. Analysis settings are listed in Table S2.

#### Figure S8

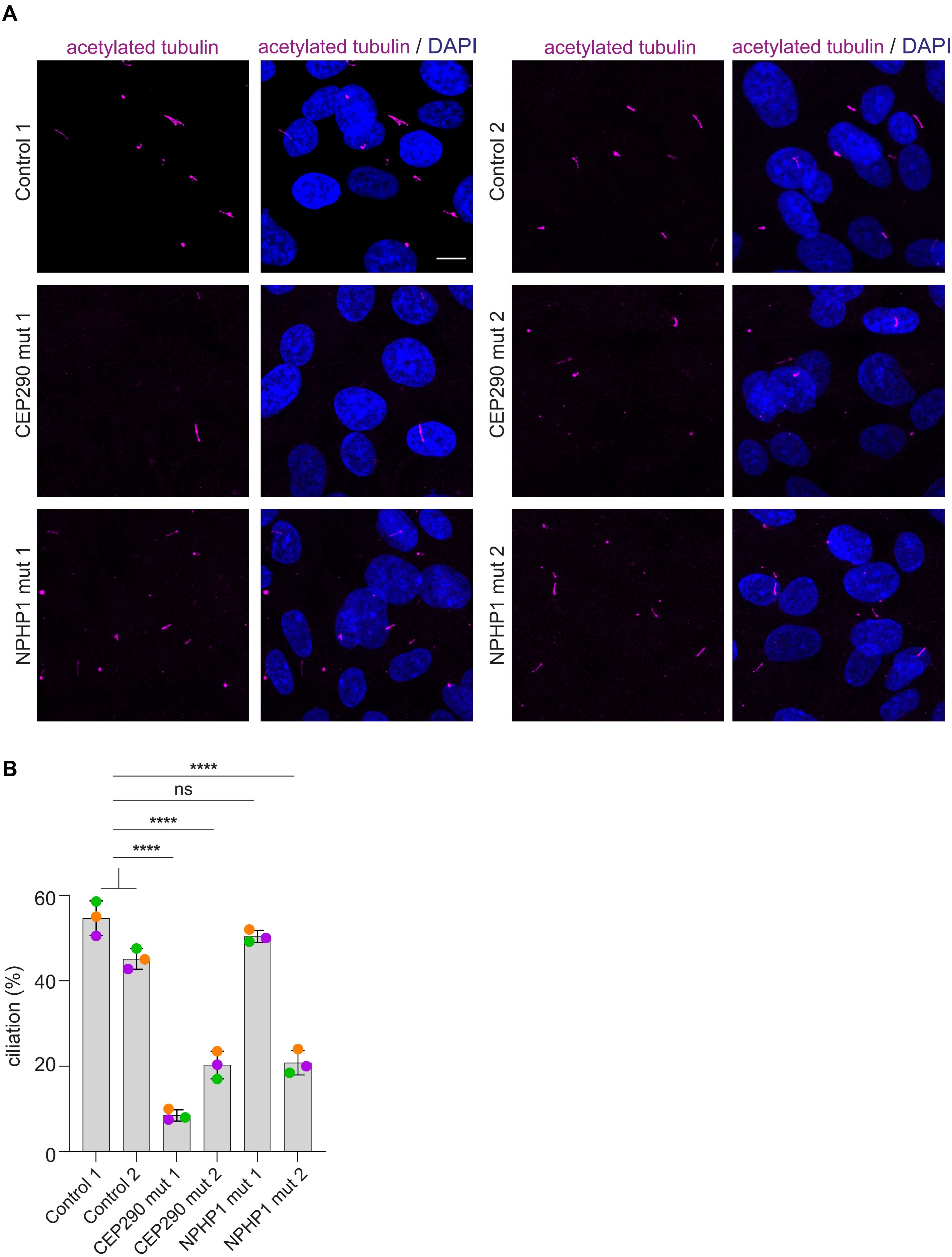
A
acetylated tubulin
acetylated tubulin / DAPI
acetylated tubulin
acetylated tubulin / DAPI
Control 1
Control 2
CEP290 mut 1
CEP290 mut 2
NPHP1 mut 1
NPHP1 mut 2
B
****
ns
****
****
ciliation (%)
60
40
20
0
Control 1
Control 2
CEP290 mut 1
CEP290 mut 2
NPHP1 mut 1
NPHP1 mut 2

**Supplementary Figure 8. Reduced ciliation in patient-derived URECs. (A)** Representative images of URECs from two healthy donors (Control 1 and 2), two patients carrying CEP290 mutations (CEP290 mut 1 and 2) and two patients carrying NPHP1 mutations (NPHP1 mut 1 and 2), ultured at the non-permissive temperature of 39 °C for 3 days to induce ciliogenesis and stained for acetylated tubulin (magenta) and DAPI (blue). **(B)** Percentage of ciliated cells for each line. Each dot represents one biological replicate; bars represent mean ± SD. Comparisons are against Control 1. Results are from 3 biological replicates (minumum 50 cells per line). Statistical significance was determined by one-way ANOVA with multiple comparisons against Control 1 and 2. Comparisons against Control 2 yielded the same statistical significance outcomes as those against Control 1, as indicated by the brackets. ****, $P < 0.0001$; ns, not significant. Scale bar: 10 µm.

#### Tables

**Table S1. Definitions and mathematical formulations of SatelliteQ measurements.** Parameters reported by SatelliteQ, with their units, mathematical definitions and the variables used in the Fiji and Python workflows. a.u., arbitrary units.

**Table S2. SatelliteQ analysis settings used for each figure.** Analysis mode, satellite rolling-ball radius, thresholding method, minimum and maximum satellite object size, watershed use, and whether centrosome and cell boundary channels were included, for every quantification shown in this study. Object sizes are given in µm² for 2D analyses and in voxels for 3D analyses. Within each experiment, identical settings were applied to all images and conditions. For Fig. S4, the settings were varied intentionally as indicated in the figure.

**Table S3. Antibodies, chemicals and software used in this study.** Primary antibodies with species, source, identifier and dilution; chemicals and reagents with supplier and identifier; and software with versions.
